# Pauperization of Emerin from nuclear envelope during chromatin bridge resolution drives prostate cancer cell migration and invasiveness

**DOI:** 10.1101/2023.11.06.565767

**Authors:** Marta Popęda, Kamil Kowalski, Tomasz Wenta, Galina V. Beznoussenko, Michał Rychłowski, Alexander Mironov, Zeno Lavagnino, Sara Barozzi, Julia Richert, Rebecca Bertolio, Jolanta Szade, Kevin Miszewski, Marcin Matuszewski, Anna J. Żaczek, Luca Braga, Giannino Del Sal, Natalia Bednarz-Knoll, Paolo Maiuri, Paulina Nastały

## Abstract

Micronuclei (MN) can arise from many causes, including the breakage of aberrant cytokinetic chromatin bridge. The frequent observation of MN in tumors raises the specter that they might not merely be passive elements but could instead play active roles in tumor progression. Here, we test the hypothesis that the presence of micronuclei could induce specific phenotypic and functional changes to the cell and lead to increased cancer invasive potential. With a variety of imaging and molecular methods *in vitro* and in clinical samples from prostate cancer (PCa) patients, we show that chromosome bridge resolution can lead to EMD accumulation and formation of EMD-rich MN. Such structure is negative for Lamin A/C and positive for LBR and Sec6β. It can cause EMD pauperization from NE affecting migratory and invasive properties of a cell and can be translated to PCa patient’s poor prognosis.

## Introduction

Prostate cancer (PCa) is the second cancer occurring in men, remains a leading cause of death in men worldwide (Sung *et al*, 2021). Chromosomal instability in PCa has been linked with higher Gleason score (Timms *et al*, 2016) and increased metastatic potential, leading to higher lethality (Miller *et al*, 2020; Stopsack *et al*, 2019; Dhital *et al*, 2023). Micronuclei (MN), one of the markers of chromosomal instability, can be defined as small, chromatin-containing nuclear structures that are physically separated from the main nucleus of a cell (Hatch *et al*, 2013). Their structure and nuclear envelope (NE) composition significantly differ from the primary nucleus (Hatch *et al*, 2013; Liu *et al*, 2018; Maass *et al*, 2018). They can arise from lagging chromosomes or chromosome fragments caused by mitotic errors or DNA damage coupled with chromatin bridge breakage (Cimini *et al*, 2003; Fenech *et al*, 2011). Micronuclei have long been used as biomarkers of genotoxicity, tumor risk, and tumor grade (Stich & Rosin, 1984; Thompson & Compton, 2011; Fenech *et al*, 2011). However, the frequent observation of micronuclei in tumors raises the specter that they might not merely be passive warnings but could instead play active roles in tumor progression e.g by inducing chromothripsis (Liu *et al*, 2018) or by activating innate immune signaling pathways (Mackenzie *et al*, 2017; Flynn *et al*, 2021). It has never been shown if presence of micronuclei could induce phenotypic and functional changes to the cell and lead to its increased invasive potential. Recent studies suggest that the composition of proteins at the NE is not even throughout the whole membrane and can show zones with accumulation of some proteins, e.g. inner and outer nuclear membrane protein, Emerin (EMD) in relation to cells’ polarization (Procter *et al*, 2020; Nastały *et al*, 2020). In the current study, we propose that accumulation of EMD in the micronucleus can lead to its consequent pauperization at the NE, increasing tumor cell migration and invasiveness in prostate cancer.

## Results

### EMD-rich MN are prevalent in prostate cancer and correlate with worse prognosis

Emerin (EMD), an inner and outer nuclear membrane protein (Nastały *et al*, 2020), has been reported as one of the “core” NE protein found in micronuclei, and on lagging chromosomes (Liu *et al*, 2018; Maass *et al*, 2018). Intriguingly, a variety of prostate cancer cell lines (LnCAP, DU145 and PC-3) often harbored micronuclei that stained extremely bright for EMD and that were very rarely found in normal prostate cell line, RWPE-1(Fig. 1A). In such structures, the EMD staining intensity exceeded nearly twice the one at the NE (Extended Data Fig. 1A), where the protein is expected to localize. We further referred to these structures as “EMD-rich MN”. The presence of EMD-rich MN was not associated with differences in EMD total protein levels in these cell lines (Extended Data Fig. 1B). To test whether EMD-rich MN are genuine cancer-associated structures or instead are an artifact of some cancer cell-lines, we then performed immunofluorescent staining of prostate tissue. Confirming our results in cancer cell-lines, we found very bright, EMD-rich structures (0-0.4315, 25^th^ to 75^th^ 0.07538-0.1861, median:0.1280, Extended Data Fig. 1C) with much higher frequency in prostate cancer (PCa) comparing to benign prostatic hyperplasia (BPH) and normal prostate tissue (Fig. 1B). Most importantly, higher numbers of EMD-rich structures were correlated to tumor higher Gleason score (prostate cancer grading system), presence of metastasis after radical prostatectomy and patients’ prognosis (expressed as time to biochemical recurrence time to Prostate-specific antigen, PSA increase after radical prostatectomy) after radical prostatectomy (≥75^th^ percentile vs <75^th^ percentile: HR: 2.64; 95% CI: 1.34-5.20; p-value Cox regression: 0.0051, Fig. 1C). No correlation to T status, tumor size and preoperative serum PSA level was found (Extended Data Fig. 1D).

**Fig. 1.**
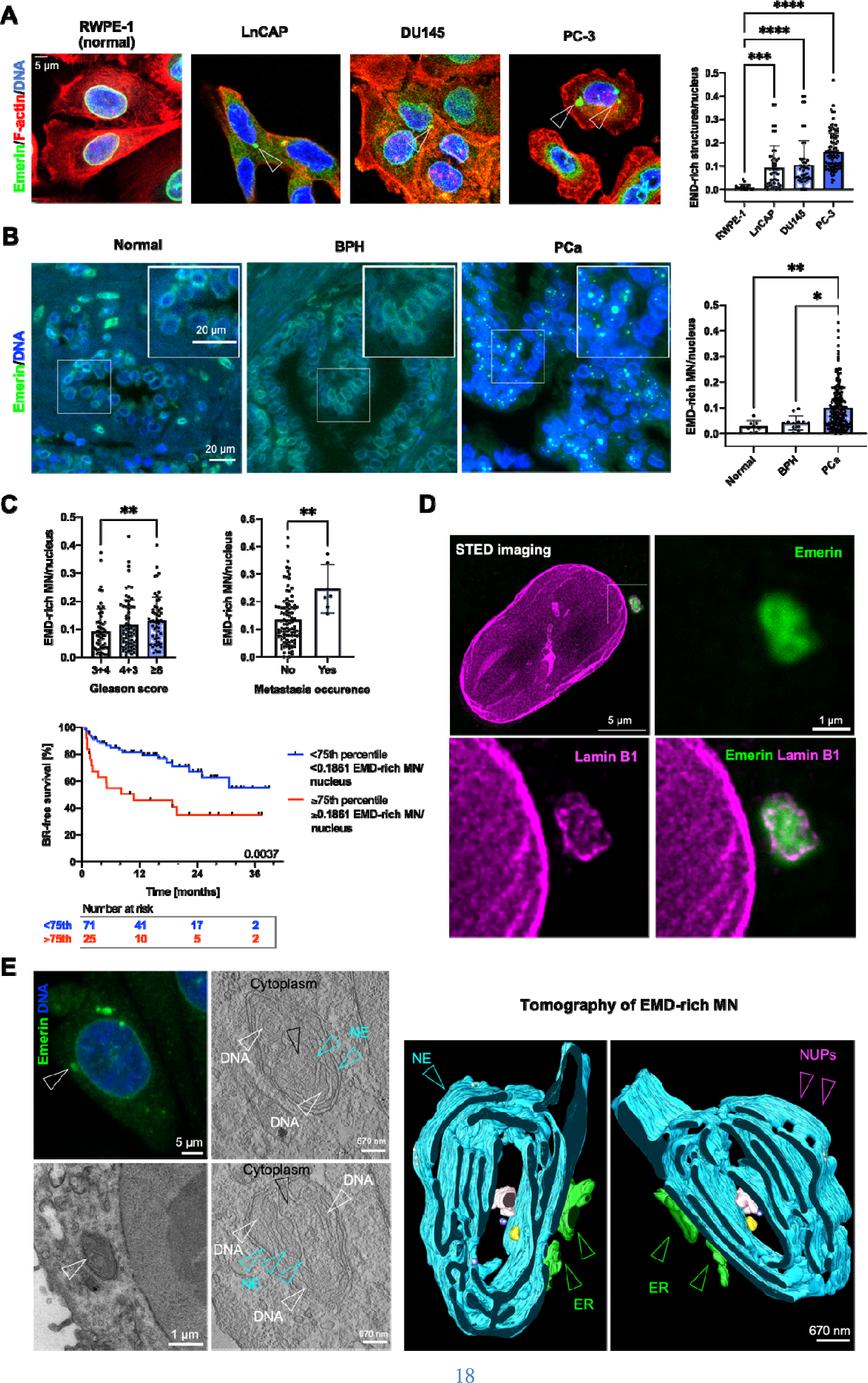
EMD-rich MNs show characteristic ultrastructure and are prevalent in PCa, where they correlate with more aggressive disease. **a** Distribution of EMD-rich MN among normal prostate epithelial cell line (RWPE-1, n=20 fields of view containing >250 cells) and prostate cancer cell lines – LnCAP (n=46 fields of view, containing >250 cells), DU145 (n=48 fields of view, containing >250 cells), PC-3 (n=98, containing >250 cells). **b** Distribution of EMD-rich in normal prostate tissue (n=4 individuals), benign prostatic hyperplasia (BPH patients), n=8, and prostate cancer (PCa, n=221 tissue cores). **c** Clinical associations between EMD-rich MN detected within primary tumor and Gleason score (n=180 tissue cores), development of metastasis after prostatectomy (n=107), and time-to biochemical recurrence (n=97 patients). **d** Super Resolution Stimulated Emission Depletion (STED) microscopy of EMD-rich MN stained for EMD and Lamin B1, EMD is shown in green, Lamin B1 in magenta. **e** Correlative light electron microscopy (CLEM) of EMD-rich MN. Upper left panel fluorescence confocal microphotography and associated electron microscopy. Right panel presents tomography of EMD-rich MN, NUPs-nuclear pores. Kaplan-Meier plots: Log-rank test; Error bars indicate 95%CI. Scatter-plot with bar (mean with SD): Mann-Whitney U test, for multiple comparison Kruskal-Wallis test was used. Error bars indicate SD. *P < 0.05; **P < 0.01; ***P < 0.001; ****P < 0.0001; ns, not significant. Source data is provided as Source data file.

To investigate if such pattern of staining could be due to EMD mutations or gene expression changes we interrogated public repositories (The Cancer Genome Atlas and cBio Portal for Cancer Genomics). We found that *EMD* gene is very rarely mutated (<0.4% out of 1,607 cases of prostate adenocarcinoma) and its gene expression can slightly increase with Gleason score (Extended Data Fig. 1E). Its expression is also slightly higher in castration-resistant prostate cancer when compared to primary tumors (Bolis *et al*, 2021) (Extended Data Fig. 1E). Thus, since EMD deregulation is associated to higher metastatic potential (Reis-Sobreiro *et al*, 2018; Liddane *et al*, 2021; Liddane & Holaska, 2021) but no frequent mutations or significant genes expression changes are found in cancer patients, alternative ways to modulate EMD presence at the NE might exist, e.g. through its aberrant localization (Reis-Sobreiro *et al*, 2018).

### EMD-rich MN have distorted NE membranes

We then investigated the fine structure of EMD-rich MN with super resolution microscopy (STED) and with correlative light electron microscopy (CLEM). Using STED, we observed that such EMD-rich MN contained disrupted sheets of lamin B1 (Fig. 1D, Extended Data Fig. 2A). Then, CLEM experiment revealed that EMD-rich MN are composed of extensive NE membrane distortions with attached chromatin, often with adjacent ER and cytoplasm protruding to the inside of these structures (Fig. 1E, Extended Data. Fig. 2B). Immunogold electron microscopy further confirmed that EMD presence was tightly associated with deformed nuclear membranes (Extended Data Fig. 2B). Interestingly, such structure of EMD-rich MN, with the clear pattern of distorted membranes resembled the one of ruptured micronuclei (Hatch *et al*, 2013; Vietri *et al*, 2020).

**Fig. 2.**
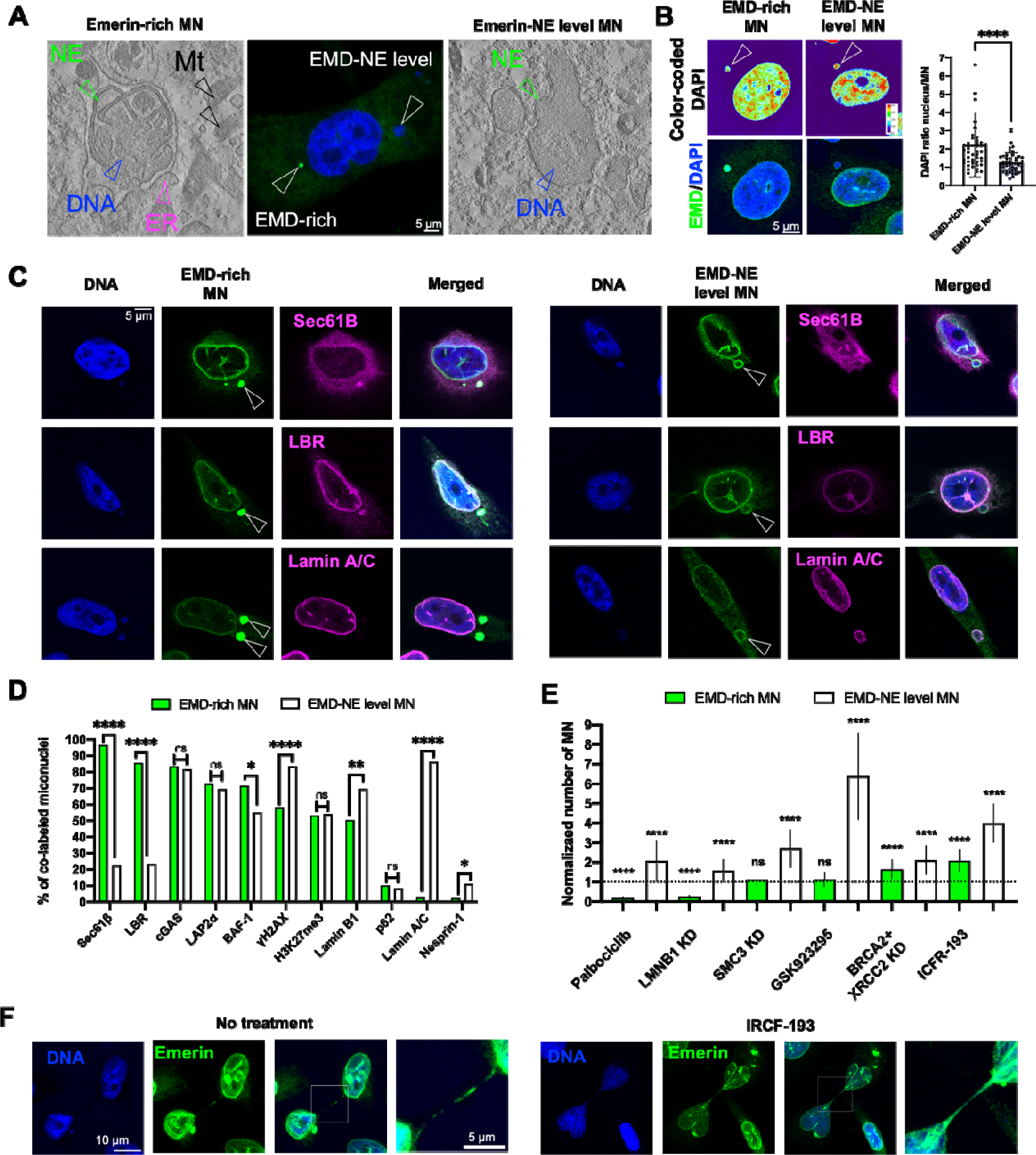
EMD expression level can differentiate micronuclei. **a** Comparison between EMD-rich MN and EMD-NE-level MN using correlative light electron microscopy (CLEM) of EMD-rich MN. **b** DAPI signal ratio quantification in EMD-rich MN (n=51 cells) and EMD-NE-level MN (n=50 cells). **c** Representative micrographs of staining for Sec6β, Lamin B receptor (LBR) and Lamin A/C in EMD-rich MN and EMD-NE-level MN, EMD is presented in green, other proteins in magenta. **d** Quantification of percent of EMD co-labeling in two MN fractions, minimally 250 cells were quantified for each condition. **e** Quantification of EMD-rich and EMD-NE level micronuclei fractions occurrence after gene silencing and treatment with selected drugs. KD-knock-down, minimally 250 cells were quantified for each condition. **f** Representative micrographs of EMD staining in chromatin bridges without treatment and after treatment with IRCF-193 inhibitor, EM is presented in green. Proportions quantified using Fisher’s Exact test are presented in bar plot. Scatter-plot with bar (mean with SD) and bar-plots Mann-Whitney U test. Error bars indicate SD.*P < 0.05; **P < 0.01; ***P < 0.001; ****P < 0.0001; ns, not significant. Source data is provided as Source data file.

### EMD-rich MN differ in composition from MN with EMD-NE levels

To specifically characterize EMD-rich MN, we investigated their composition using multiple markers. We first classified micronuclei by their EMD intensity level, higher than NE or at NE-level, and we then compared their ultrastructure. Interestingly, MN with EMD-NE level did not show the same pattern of NE membrane distortions as observed in EMD-rich ones (Fig. 2A, Extended Data Fig. 3A). Instead, EMD-rich MN showed a decreased DAPI signal, a good proxy of DNA content (20), and an increase in area, probably due to their excess of membranes (Extended Data Fig. 3B). M**Error! Bookmark not defined.**icronuclei typically present defective NE that can result in enrichment in core NE proteins (EMD, LAP2α, BAF, Lamin A/C) or in non-core ones, such as nucleoporins, Lamin B1, Lamin B receptor (Liu *et al*, 2018; Dechat *et al*, 2004; Kwon *et al*, 2020). Therefore, we tested if EMD-rich MN might differ in composition from micronuclei with EMD-NE level. As expected, for their high membrane content, EMD-rich MN co-labeled more often with endomembranes marker, Sec6β (Ferrandiz *et al*, 2022), and EMD interactor, BAF-1 (Berk *et al*, 2014) (Fig. 2C-D and Extended Data Fig. 3C). Contrary, EMD-NE MN were higher in DNA damage marker γH2AX (Hatch *et al*, 2013) and lower in Lamin B1 (1,21). cGAS, that localize to MN upon nuclear envelope rupture (Mackenzie *et al*, 2017), was found in ∼80% of both MN fractions. Emerin-rich MN were also more positive for Lamin B receptor (LBR), and mostly negative for and Lamin A/C, when compared to EMD-NE level ones (Fig. 2C-D). Lamin A/C, was previously shown to mark MN nearly in 100%, which was true for EMD-NE level MN, as showed in Fig.2C-D (Kneissig *et al*, 2019). Surprisingly, another LEM-domain containing protein, LAP2α (Brachner & Foisner, 2014), was found with the same occurrence in both MN types (Fig.2C-D). Similarly, a lysine trimethylation on histone H3 (H3K27me3), often enriched in MN (Agustinus *et al*, 2023), was also found with the same frequency in both MN types. Taken together, the collected data suggested that EMD level can differentiate MN into two fractions. So, we further asked a question whether the sub-type of EMD-rich MN could form from a different event that other fractions of MN.

### EMD-rich MN originate during chromatin bridge resolution

We tested various known MN-inducing and affecting conditions to see if they can promote one or the other fraction of MN. The depletion of lamin B1, that is known to trigger NE disruption (Hatch *et al*, 2013), induced a decrease in the frequency of EMD-rich MN and an increase in the one of EMD-NE level (Fig. 2E). Interestingly, EMD-NE level MN occurrence exclusively increased after reported MN-inducting conditions including: SMC3 cohesin knockdown (Leylek *et al*, 2020), treatment with CENP-E inhibitor, GSK923295 (Ferrandiz *et al*, 2022), and palbociclib incubation (Crozier *et al*, 2022) (Fig. 2E and Extended Data Fig 4A-B). Whereas conditions as silencing genes involved in DNA repair (*BRCA2* and *XRCC2*) and topoisomerase II inhibitor, IRCF-193 generally increased the frequency of both kind of MN (Fig. 2E and Extended Data Fig 4A-B). As expected, all treatments generally increased the number of both type of MNs. However, interestingly, the conditions that more specifically induced EMD-rich versus EMD-NE MN were the ones known to increase the formation of aberrant chromatin bridge upon failed abscission: the simultaneous knock down of *BRCA2* and *XRCC2* (Liu *et al*, 2018; Flynn *et al*, 2021; Umbreit *et al*, 2020; Jiang *et al*) or the topoisomerase inhibitor, IRCF-193 (30) (Fig. 2E and Extended Data Fig 4A-B). This results therefore suggested that EMD-rich MN can originate from collapsed or miss-resolved chromatin bridges.

To test this hypothesis, we performed timelapse analysis of a cell cycle in cells transfected with EMD-EGFP construct and treated with IRCF-193. Chromatin bridges are aberrant structures occasionally occurring also in untreated cells and that were showed to persist for 3-20h (Maciejowski *et al*, 2015), while in our case they lasted for 1.2 – 14.7 h with the median of ∼6h. We confirmed presence of EMD in these structures, and moreover, our data strongly suggest that EMD-rich MN were formed during the resolution of a chromatin bridge (Fig. 3A, Supplementary Movie 1 and 2). Out of all imaged cell divisions, EMD-rich MN formation was the most frequent event associated with chromatin bridge resolution occurring in at ∼35% of investigated events (Fig. 3B). EMD-rich MN formation was associated with longer chromatin bridge formation and resolution (Fig. 2C), whereas shorter chromatin bridges led often to binucleated cell formation (Fig. 3A-B, Supplementary Movie 3). We did not observe correlation with EMD-rich MN formation and time of chromatin bridge persistence (Fig. 3D). Interestingly, we also observed that appearance of EMD-rich MN changed completely this proteins’ NE to cytoplasm ration (Fig. 3E), which indicated that EMD could be pauperized from NE in favor of micronucleus.

**Fig. 3.**
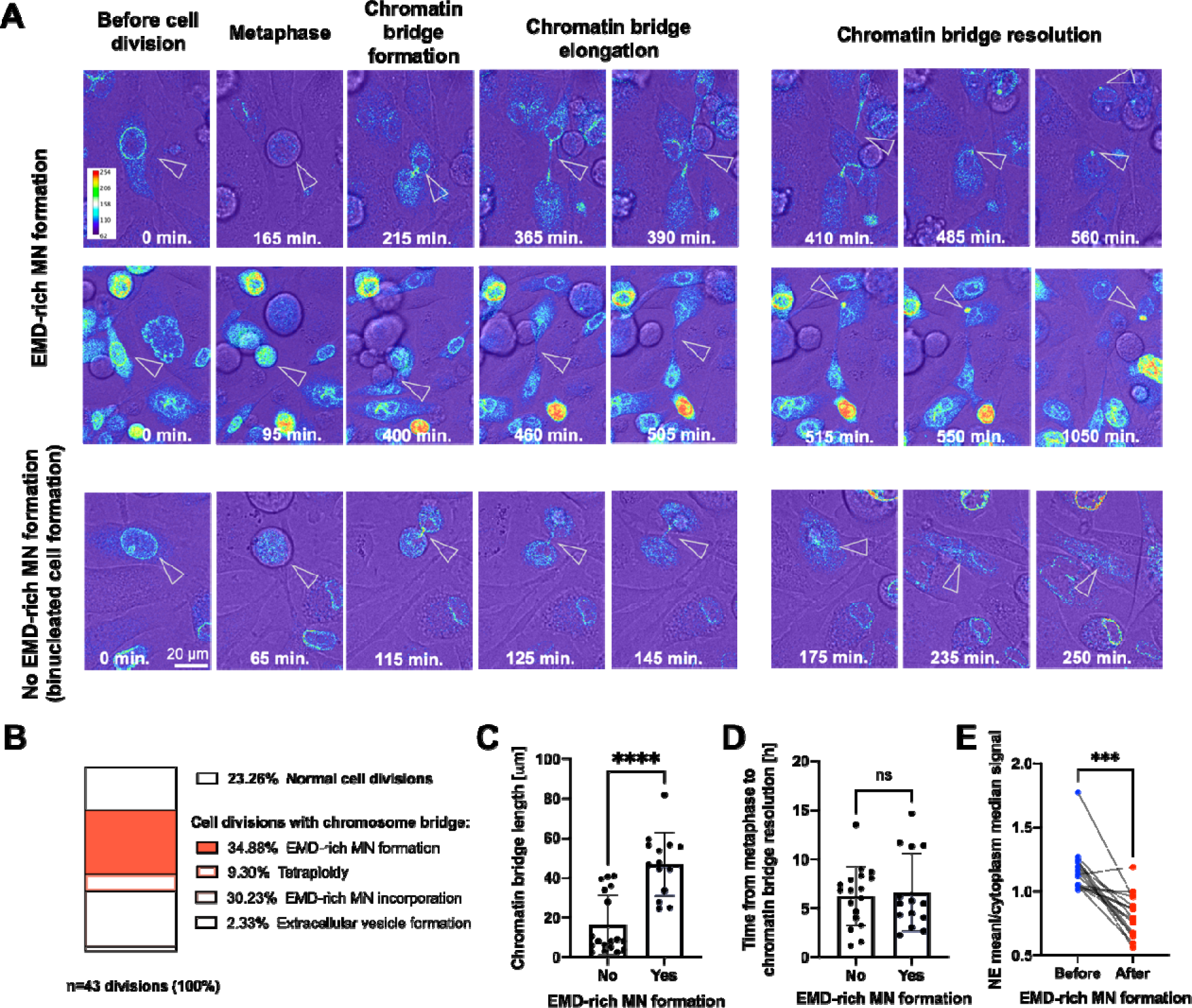
EMD-rich MN are associated with chromatin bridge resolution. **a** Representative micrographs of cell divisions after IRCF-193 treatment of PC-3 cells transfected with EMD-EGFP construct. Arrows indicate chromatin bridges and EMD-rich MN. **b** Percentages of events associated with cell division in all imaged cells. **c** Quantification of chromatin bridge length in cell divisions with (n=14) and without (n=18) EMD-rich MN formation. **d** Quantification of chromatin bridge persistence in cell divisions with (n=14) and without (n=18) EMD-rich MN formation. **e** Quantification of NE mean/cytoplasm median of EMD signal before and after EMD-rich MN formation (n=15). Scatter-plot with bar (mean with SD): Mann-Whitney U test was used. Paired analysis: Wilcoxon matched-pairs signed rank test. *P < 0.05; **P < 0.01; ***P < 0.001; ****P < 0.0001; ns, not significant. Source data is provided as Source data file.

### EMD-rich MN cause EMD pauperization from NE

In general, decreased EMD expression during tumor transformation is associated with the nuclear structural defects required for increased cancer cell migration and invasiveness, leading to increase metastatic potential and unfavorable prognosis (Liddane & Holaska, 2021; Reis-Sobreiro *et al*, 2018; Liddane *et al*, 2021). As EMD accumulation was so evident in EMD-rich MN, we observed that cells with EMD-rich MN showed pauperization of this protein from NE (Fig. 4A). Emerin, in addition to its localization at the inner and outer nuclear membrane, is also present at the ER membranes (Nastały *et al*, 2020; Buchwalter *et al*, 2019), which could explain the accumulation of membranes and EMD in the EMD-rich MN. To address if there is interconnection between EMD-rich MN and NE, we performed an iFRAP experiment (inverse fluorescence recovery after photobleaching). Emerin was photobleached from NE in EMD-GFP-transfected cells, which was associated with fluorescence signal decay from EMD-rich micronuclei (Fig. 4B, Supplementary Movie 4. Similarly, when EMD-rich MN was photobleached, EMD-GFP signal intensity decreased at the NE (Fig. 4C, Supplementary Movie 5). Additionally, the pattern of EMD traffic resembled vesicle-based traffic, rather than diffusion (Beznoussenko *et al*, 2014), which additionally confirms EMD association with the membranes. The collected data showed that EMD can shuffle between these two compartments, which could be a way for the cell to modulate its presence in both MN, and NE. As we confirmed that NE pauperization can be achieved, we further aimed to test if it can be also translated to cellular phenotype.

**Fig. 4.**
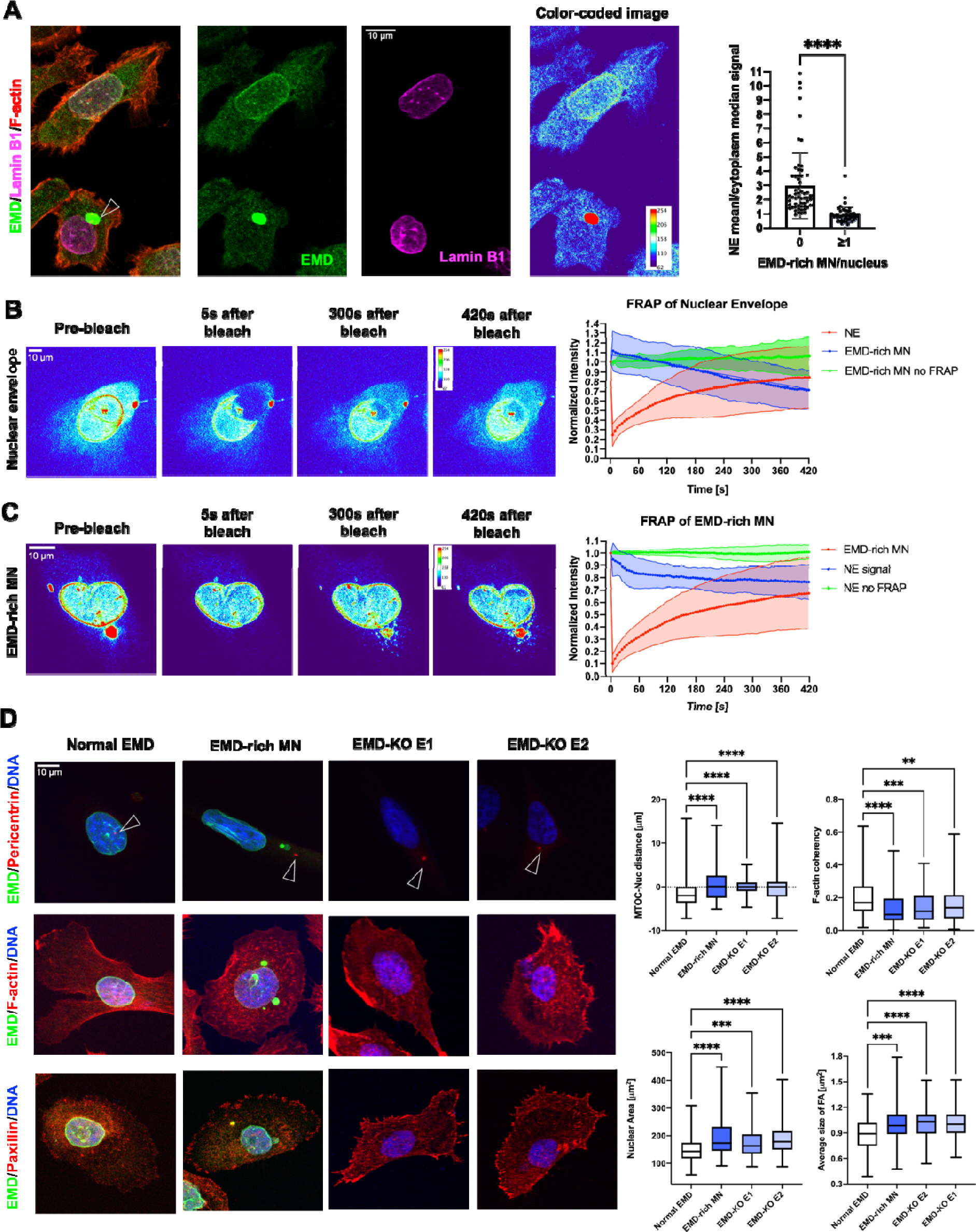
EMD-rich MN can induce emerin pauperization form NE and lead to EMD-loss phenotype recapitulation. **a** Quantification of NE mean/cytoplasm median of EMD signal in fixed PC-3 cells with (n=51) and without EMD-rich MN (n=62). **b** iFRAP analysis of PC-3 cells transfected with EMD-EGFP construct. The region of nuclear envelope was bleached (n=23 cells), control without bleaching (n=20 cells). **c** iFRAP analysis of PC-3 cells transfected with EMD-EGFP construct. The region of EMD-rich MN was bleached (n=15 cells), control without bleaching (n=20 cells). **d** Quantification of selected parameters in PC-3 cells without EMD-rich MN, with EMD-rich MN, EMD-KO E1 and EMD-KO E2: nucleus-MTOC distance (n=123, n=52, n=79, n=106, respectively), F-actin coherency (n=145, n=66, n=136, n=167, respectively), nuclear area (n=123, n=52, n=79, n=106, respectively), and focal adhesion size (n=221, n=73, n=130, n=141, respectively). Scatter-plot with bar (mean with SD): Mann-Whitney U test. Box-plot; Whiskers indicate min to max values, within the box the first quartile, median, third quartile are represented. For multiple comparison Kruskal-Wallis test was used. *P < 0.05; **P < 0.01; ***P < 0.001; ****P < 0.0001; ns, not significant. Source data is provided as Source data file.

### Presence of EMD-rich MN recapitulates EMD loss at the cellular level

Emerin loss is associated with characteristic cellular phenotype that includes increased distance between MTOC and nucleus (Nastały *et al*, 2020; Chang *et al*, 2013; Hale *et al*, 2008), more random F-actin fiber organization due to altered actin flow (Chang *et al*, 2013), alterations in focal adhesion size (Nastały *et al*, 2020), increased nuclear area and irregular shape of the nucleus (Lammerding *et al*, 2005), and increased migratory and invasive properties (Nastały *et al*, 2020; Reis-Sobreiro *et al*, 2018; Liddane *et al*, 2021). Emerin decreased expression and deregulation has been also reported to increase metastatic potential of prostate cancer cells (Reis-Sobreiro *et al*, 2018). To address the question if presence of EMD-rich MN could recapitulate the EMD-loos phenotype, we compared some of the phenotypic changes in cells without MN, cells with EMD-rich MN and EMD knock-out (EMD-KO) cells (Extended Data Fig. 5A). Indeed, cells containing EMD-rich MN resembled phenotypically cells with EMD deficiency (Fig. 3C), suggesting that accumulation of EMD in the cytoplasmic structure can lead to altered properties of the cell. Importantly, cells with EMD deficiency phenotype caused by presence of EMD-rich MN or EMD-KO had larger size of focal adhesion, suggesting more migratory phenotype (Schmidt *et al*, 2022; Kim & Wirtz, 2013). We, therefore, further investigated the cells with presence of EMD-rich MN and EMD pauperization signature in context of prostate cancer.

### The transcriptomic signature of pauperized EMD in PCa

To verify the signature of the presence of EMD-rich MN or EMD loss in prostate tumors, their transcriptomic profile was analyzed using nCounter PanCancer Progression Panel. Based on immunofluorescent staining for EMD, tumors were divided in two groups: normal EMD NE staining and pauperized EMD from NE (presence of EMD-rich MN or EMD loss in cancer cells, Fig 5A). Tumors with EMD pauperized form NE had up-regulated 33 genes (Supplementary Table S4) including the ones encoding proteins involved in prostate cancer migration and invasion as vimentin (Lang *et al*, 2002), SPARC (López-Moncada *et al*, 2022), or invasion as MMP2 (Trudel *et al*, 2003), as presented in Fig 5A and Extended Fig. 5B. The gene ontology analysis revealed that among processes enriched in tumors with pauperized EMD were cell adhesion, positive regulation of cell migration and response to TGFβ pathway (Fig. 5B). When we grew EMD-KO PC-3 cells in 3D culture conditions, we observed that they had more invasive morphology – the 3D spheroids were larger, more branched, and less circular when comparing to the ones form the control cells (Fig. 5C, Extended Data Fig. 5C). We further analyzed them using RNA sequencing (Extended Data File 1). Then, we verified if PC-3 cells grown in 3D condition (EMD-KO) could share any transcriptomic signature with the tumor cells form with pauperized EMD from PCa. Indeed, there was a common pattern of gene upregulation in both datasets (Fig. 5D). The top 9 up-regulated genes in both datasets included *CXCR4*, *APOE*, *SPARC*, *VIM*, *GSN*, *ANXA2P2*, *SFRP1*, *COL18A1*, *FN1* (Fig 5D, Supplementary Table S5).

**Fig. 5.**
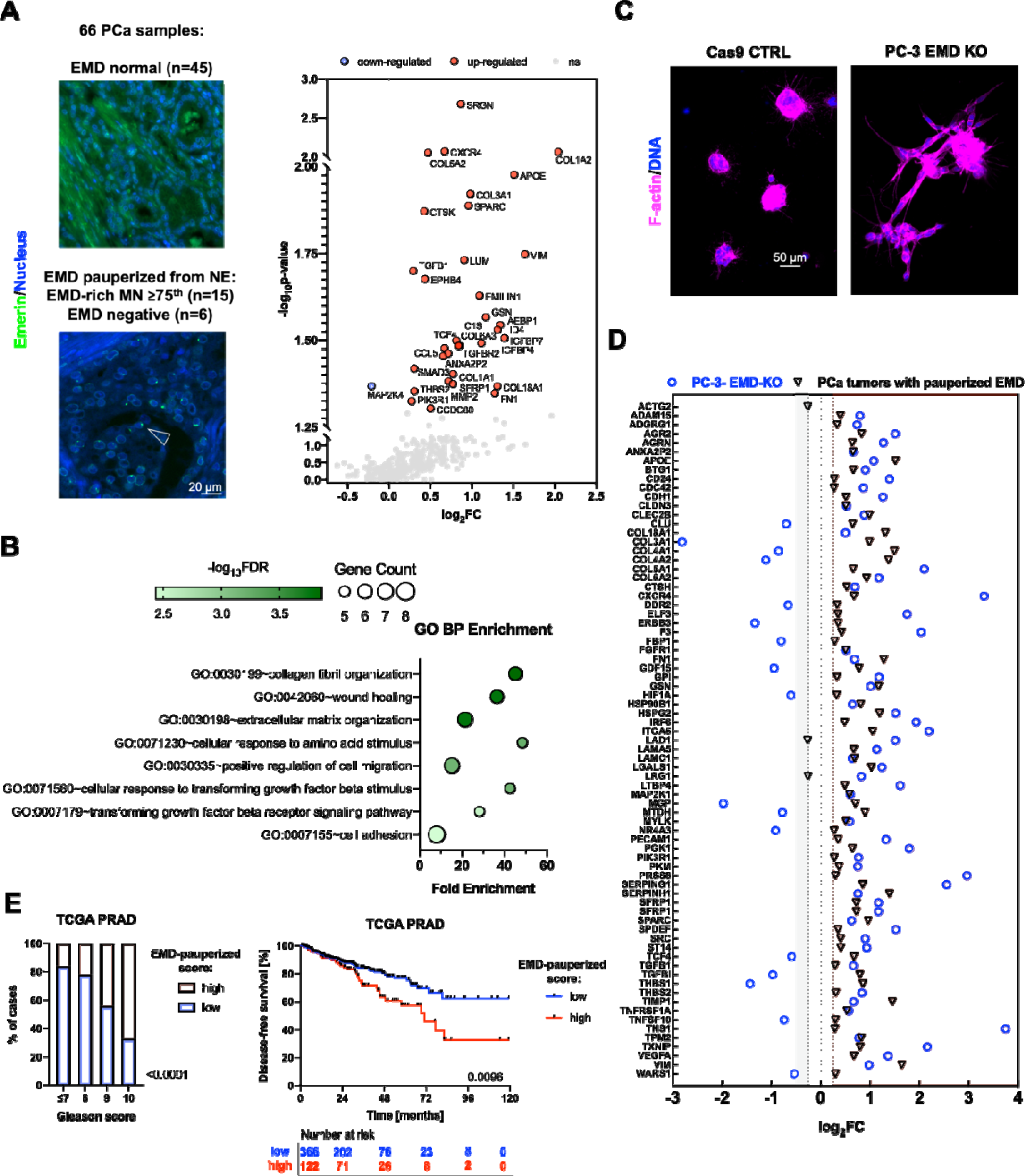
The transcriptomic signature of pauperized EMD in PCa. **a** nCounter analysis of prostate tumors (n=66) divided into 2 groups based on EMD: normal NE expression of EMD and EMD pauperized from NE (presence of EMD-rich MN or EMD negative). Scatter plot representing differentially expressed genes. **b** Gene ontology analysis of up-regulated genes in tumors with pauperized EMD at the NE. **c** PC-3 cells with EMD-KO growth in 3D culture conditions **d** Forest plot of 75 genes that were common in both datasets – the nCounter Nanostring dataset for prostate tumors and RNA sequencing dataset from PC-3 cells grown in 3D conditions. To facilitate visual representation *ACTG2* gene point for EMD-KO group is not presented at the graph (log_2_FC= 5.861). **e** The application of EMD-pauperized in TCGA PRAD dataset. Its association to Gleason score and disease-free survival prediction (n=488). Bar plot: Chi-square test. Kaplan-Meier plot: Log-rank test; Error bars indicate 95%CI. P < 0.05; **P < 0.01; ***P < 0.001; ****P < 0.0001; ns, not significant. Source data is provided as Source data file.

### EMD-pauperized transcriptomic score predicts shorter time to progression in TCGA dataset

Eventually, we verified the potential adverse impact of EMD pauperization on prostate cancer prognosis in TCGA dataset. We computed transcriptomic score including top 9 up-regulated genes in prostate tumor tissues and EMD-KO experiments (*CXCR4, APOE, SPARC, VIM, GSN, ANXA2P2, SFRP1, COL18A1, FN1*) and confronted it with patients’ characteristics and disease-free survival (DFS) defined as time to disease recurrence or progression. The EMD-pauperized score increased with Gleason score and importantly, high score was linked to shorter time to DSF (HR: 1.75; 95% CI 1.14-2.69; p-value: 0.0106 as presented in Fig. 5E). The collected data from the tumor samples collected from PCa patients and 3D PC-3 cell spheroid culture suggests that cells with EMD pauperization share increased migratory and invasive properties that could drive their metastatic potential and indicate poor prognosis in PCa patients. We further aimed to verify if there is an enrichment of cells with EMD-rich MN in motile and invasive cells using functional assays.

### Cells with EMD-rich MN show increased migratory and invasive potential

To functionally test if cells with EMD-rich MN are characterized with migratory phenotype, we firstly performed 2D random migration analysis. We observed that cells with EMD-rich structures tend to move faster and for longer pathways (Fig. 6A), which was previously observed in EMD-deficient cells (Nastały *et al*, 2020). Similarly, within the transwell assay, cells that migrated through constriction had significantly more EMD-rich MN comparing to the control cells (Fig. 6B). The frequency of cells with EMD-rich MN was much higher in invading cells, comparing to the non-invading ones as observed in spheroid culture (Fig. 6C) and collagen-invasion assay (Fig. 6D). Taken together our data suggests that the formation of EMD-rich MNs, inducing a functional pauperization of EMD at the nuclear envelope, causes a specific phenotype switch in prostate cancer cells, finally inducing a more invasive behavior strictly linked to a worst prognosis.

**Fig. 6.**
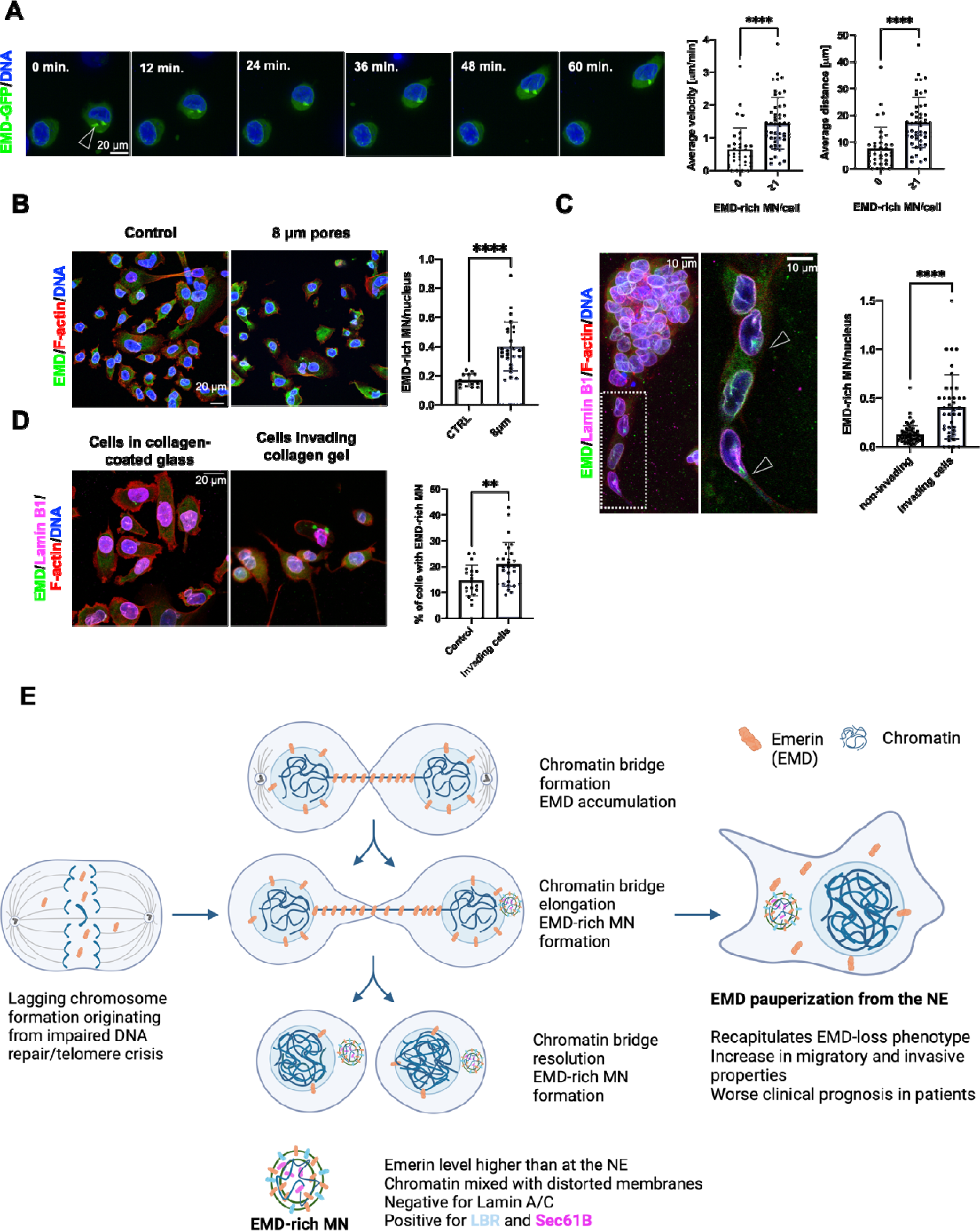
Cells with EMD-rich MN show increased migratory and invasive properties. **a** Assessment of migratory potential of cells with (n=50) and without (n=36) EMD-rich MN in PC-3 cells transfected with EGFP-EMD. Average cells’ velocity and distance were calculated from individual paths. **b** Quantification of EMD-rich MN/nucleus in PC-3 cells in control conditions (n=13 fields of view with >250 cells) and in cells that in passed 8 µm constriction (n=25 fields of view with >250 cells). **c** Assessment of EMD-rich MN/nucleus in non-invading (n= 55 spheroids) and invading cells (n=43 spheroids) from PC-3 cultured in 3D. **d** Quantification of EMD-rich MN/nucleus in PC-3 cells in control conditions (n=19 fields of view with >250 cells) and in cells that invaded collagen (n=28 fields of view with >250 cells) in collagen-invasion assay. **e** A proposed mechanism of EMD-rich formation during chromatin bridge resolution followed by phenotypic change. Scatter-plot with bar (mean with SD): Mann-Whitney U test. *P < 0.05; **P < 0.01; ***P < 0.001; ****P < 0.0001; ns, not significant. Source data is provided as Source data file.

## Discussion

In the current study, we showed that presence of micronuclei could induce phenotypic and functional changes to the cell and lead to its increased migratory and invasive potential. As a mechanism we proposed that MN occurrence could cause NE pauperization of EMD protein.

Emerin mis-localization has been already reported in PCa, in 80 cases, where it discriminated cancer from benign tissue and correlated with disease progression (Reis-Sobreiro *et al*, 2018). We confirmed this observation in a slightly larger cohort of patients with increased risk of developing metastasis (defined as d’Amico intermediate to high-risk PCa) and showed that we can even stratify patients by the number of EMD-rich MN, which correlates to Gleason score, shorter time to biochemical recurrence and occurrence of metastasis after radical prostatectomy. With various imaging methods, we were able to investigate its ultrastructure and composition. We also observed that EMD amount can differentiate MN but it did not follow the previously proposed classification to “core” and “non-core” proteins (Liu *et al*, 2018) or based on the presence of rupture (Hatch *et al*, 2013; Mackenzie *et al*, 2017), suggesting their different derivation. Essentially, we found that the ultrastructure and composition of the EMD-rich MN differ from the ones of EMD-NE MN, suggesting that the two could originate from different mechanisms.

Emerin has been previously found to assemble on lagging chromosomes or chromatin bridges, with lack of LBR on these structures (Liu *et al*, 2018). Following this observation, here, we show that EMD-rich micronuclei probably form during chromatin bridge collapse. As it was observed before, indeed, chromosome bridge formation predisposes to micronucleation (Umbreit *et al*, 2020; Crasta *et al*, 2012). Moreover, we observed that EMD-rich MN formed more often in longer chromatin bridges, regardless of its lifetime and it has already been proposed that cellular mechanical forces could stretch a chromatin bridge We then hypothesize that long chromatin bridges could drive severe NE membrane accumulation, that upon the abrupt resolution of the bridge leads to the formation of micronuclei with the characteristic membrane distortions observed in EMD-rich MN. **Error! Bookmark not defined.**Such long chromatin bridge resolution, inducing EMD-rich MN formation, finally causes EMD pauperization from the NE. Emerin, indeed, can shuffle between EMD-rich MN and NE and the presence of an abundant bunch of membrane in the EMD-rich MN acts as a sink for EMD and causes its pauperization from the NE. This mechanism modulates EMD abundance at the NE, without affecting *EMD* gene expression. EMD levels at the NE regulate important cellular process, like cell’s motility (Lavenus *et al*, 2022) and cells’ polarization (Procter *et al*, 2020; Nastały *et al*, 2020). Moreover, emerin-loss, is associated with altered nuclear envelope elasticity that contributes to increased nuclear fragility (Rowat *et al*, 2006). More generally, decreased EMD expression during tumor transformation is associated with nuclear structural defects, increased cancer cell migration and invasiveness, increased metastatic potential and unfavorable prognosis (28–30). Importantly, cells positive for EMD-rich MN partially phenocopies EMD-KO cells, and they display larger focal adhesions, suggesting more migratory capacity (Schmidt *et al*, 2022; Kim & Wirtz, 2013).

Previously, EMD mis-localization has been shown in PCa and linked to its metastatic potential. Circulating tumor cells from prostate cancer patients were shown to have mis-localized EMD and prostate cancer cell line, DU145 cells with silenced EMD formed widespread metastases more efficiently in mouse model (Reis-Sobreiro *et al*, 2018). Our data from the tumor samples collected from PCa patients and 3D PC-3 cell spheroid culture suggests that cells with EMD pauperization share common transcriptomic signature. Mostly genes involved in cell migration and invasion were up-regulated, many of them were previously linked with prostate cancer progression (Lang *et al*, 2002; López-Moncada *et al*, 2022; Trudel *et al*, 2003; Lu *et al*, 2007). We also observed that the set of genes up-regulated in cells with EMD pauperization could indicate patients with worse prognosis. We were also further able to confirm the enrichment of cells with EMD-rich MN in the population of the most migratory and invasive cells using functional assays. Our observed results in increased epithelial-mesenchymal plasticity phenotype could indicate enhanced tissue fluidification (48). Such mechanically driven transcriptional rewiring is associated with cell state alteration with the emergence of malignant traits, including epithelial-to-mesenchymal plasticity phenotypes and could explain poorer patients’ prognosis.

Emerin pauperization from the NE might participate the complex series of events that drive prostate tumor progression. Collapses chromatin bridges at the same time can generate EMD-rich MN, inducing its pauperization at the NE, and also DNA damage, leading to extreme genomic complexity and continually evolving subclonal heterogeneity (Umbreit *et al*, 2020). The result is that downstream of chromosome bridge formation, the accumulating burden of DNA breakage can easily exceed the capacity to stabilize broken chromosome ends. Therefore, complex genome evolution with subclonal heterogeneity is virtually an inevitable consequence of chromosome bridge formation, itself a common outcome of cell division defects during tumorigenesis. Both genomic instability and deficient DNA repair mechanisms have been reported in PCa, where it correlates with both worse prognosis and migratory phenotype (Miller *et al*, 2020; Stopsack *et al*, 2019; Dhital *et al*, 2023; Bednarz-Knoll *et al*, 2019). Here, we demonstrate that EMD-rich could serve as a predictor in PCa. However, such data should be verified in larger cohort. Taken together, with a variety of imaging and molecular methods, we were able to show that chromosome bridge resolution can lead to EMD accumulation and formation of EMD-rich MN. Such structure is negative for Lamin A/C and positive for LBR and Sec6β. Acting as a protein sink, it causes EMD pauperization from NE. This, in turn, is functionally linked to migratory and invasive properties of a cell and can be translated to patient’s poor prognosis (Fig. 6E). Taking into account EMD ubiquitous expression across tissues and various cancers (Liddane & Holaska, 2021; Nagano *et al*, 1996), the EMD pauperization pathogenic effect could be translated into other cancerous diseases.

## Methods

### Cell lines

Human immortalized epithelial prostate cell line RWPE-1 (CRL-3607) was purchased from ATCC (American Type Culture Collection) and cultured in Keratinocyte Serum Free Medium (Thermo Fisher #17005042) and 1% Penicillin-Streptomycin (Thermo Fisher Scientific #15140122). Human prostate cell lines PC-3 was obtained from ATCC Human prostate cell lines DU145 (HTB-81), LnCAP (CRL-1740), and PC-3 (#CRL-1435) was purchased from ATCC and were cultured in medium RPMI 1640 (Thermo Fisher Scientific #11875093) supplemented with 10% FBS (Thermo Fisher Scientific #10270-106) and 1% Penicillin-Streptomycin (Thermo Fisher Scientific #15140122). The cell lines were periodically tested for mycoplasma. All cell lines were used for no more than 5-10 passages after thawing.

### Patient samples and ethic approval

The study has been approved by the local Ethics Committee (i.e., Independent Bioethics Committee for Scientific Research at the Medical University of Gdańsk, no. NKBBN/286/2018). Written informed consent was obtained from all subjects involved in the study. A total of 351 primary tumor samples corresponding to 117 d’Amico intermediate to high PCa patients, who underwent radical prostatectomy in the Department of Urology at the Medical University of Gdańsk (Poland) in years 2018–2022, were analyzed. Tumor specimens were prepared as tumor microarrays (TMAs) that consisted of 3 tumor fragments per patient, including fragments of tumor with highest and dominant Gleason score (Nastały *et al*, 2021). All the tissue cores were assessed by experienced pathologist for the percentage of tumor cells. The clinical data obtained for the patients is presented in **Supplementary Table S1**.

### Immunofluorescence staining on cells

Selected antigens were immunolocalized in cultured cells, plated on the collagen-coated coverslips. Cells were fixed with 4% PFA for 10 min., washed with 1xPBS, permeabilized with 0.1% Triton X-100/1xPBS (Sigma-Aldrich). Non-specific binding of antibodies was blocked with 3% bovine serum albumin (BSA) in 1xPBS, at room temperature, for 1h. Then, samples were incubated overnight with primary antibodies (**Supplementary Table S2**) diluted in 3% BSA in PBS, washed with 1xPBS, and incubated at room temperature with secondary antibodies diluted at 1: 500 in 1.5 % BSA in 1xPBS. After washing with 1xPBS, cell nuclei were visualized by incubation with DAPI (Thermo Fisher Scientific #D1306). Specimens were mounted with ProLong™ Gold Antifade Mounting Medium (Thermo Fisher Scientific #P36930).

### Immunofluorescence staining in tissues

The 4-6um thick TMA sections were deparaffinized and antigens were unmasked using EnVision FLEX Target Retrival Solution Low pH (Dako Agilent) in PT link device (PT200, Dako Agilent) for 20 min. at 97°C. TMAs were then incubated overnight at 4 °C with mouse monoclonal anti-EMD in vitro diagnostic antibody (4G5 clone, Novocastra) diluted 1:100, followed with secondary antibody incubation for 60 min. The nuclei were stained with DAPI and sections were mounted in Vectashield Antifade mounting medium (Vector Laboratories).

### Quantification of EMD-rich MN in tissue and cell samples

To count the number of EMD-rich MN/nucleus a custom integrated image analysis workflow for was created, where QuPath (Bankhead *et al*, 2017) and ImageJ (Schindelin *et al*, 2012; Schneider *et al*, 2012) were combined. The position of the prostate cancer cells was selected. The tumor cell presence was verified by an experienced pathologist. AMACR(Molinié *et al*, 2004) immunostaining was used as a confirmatory stain for prostatic adenocarcinoma in conjunction with H&E morphology. The number of nuclei was counted using the cell detection tool in QuPath software (ver. 0.3.0) in the selected region. The selected region was further transferred into ImageJ (ver. 2.3.0), where individual fluorescent channels were loaded into the workflow and annotated, nuclear, and EMD images were segmented in parallel. The EMD images were segmented by conventional thresholding and converted to a binary mask using ImageJ EMD-rich MN were subsequently identified (on the basis of their size: 0.2-12 um^2^ and emerin intensity). Segmentation models were improved through an iterative training process until ≥95 concordance was achieved with manual counting. For each field of view, EMD-rich MN were identified and counted using structures using ImageJ’s Analyze Particles (https://imagej.net/imaging/particle-analysis). The identification and quantitation of EMD-rich micronuclei in cells was performed with similar workflow, just without region selection in QuPath. Each field of view was then verified by observer.

### Super resolution Stimulated Emission Depletion (STED) microscopy

For STED imaging (Hell, 2007), cells were fixed with 4%PFA, blocked in 10% normal goat serum (Thermo Fisher Scientific) and incubated for 45 min. with anti-EMD (4G5 clone, Novocastra), anti-Lamin B1 (Abcam, ab16048) antibodies. Followed by incubation with secondary antibodies: anti-mouse IgG Atto-594 (Sigma-Aldrich, 76085) and anti-rabbit-IgG-Atto-647N (Sigma-Aldrich, 40839). The specimen was then mounted in Moviol (Sigma-Aldrich). The cells were imaged with HC PL APO 100x 1,4 NA STED oil immersion objective (506478, Leica, Germany) mounted on a Leica TCS SP8-STED microscope (Leica Microsystems, Wetzlar, Germany) equipped with a tunable pulsed White Light Laser Source for confocal imaging and depletion lasers (592 nm – continuous wave- and 775 nm - pulsed).

### Correlative light electron microscopy (CLEM)

The PC-3 cells were seeded on collagen-coated 35 mm glass-bottom dishes with grid (MatTek Life Sciences) and left to attach for 48h. The cells were fixed for 5 min. in 4% PFA /0.05 % Glutaraldehyde/0.15 M HEPES (pH 7.3), followed by fixation in the same buffer 3x for 10 min. The specimen was blocked for 20 min. in 1%BSA/PBS and then permeabilized for 15 min. in 0.1% Saponin 1xPBS. The cells were incubated with primary anti-mouse EMD (4G5 clone, Novocastra) antibody for 45 min, followed by 45 min incubation secondary anti-mouse antibody conjugated with Alexa-488 (Thermo Fisher Scientific). The brighfield and fluorescent images of stained cells were immediately acquired with the inverted Leica CLSM TCS SP2 microscope, equipped with 20x (air, 0.55 NA) and 40x (air, 0.95 NA) objectives. Two-step CLEM based on the analysis of tomographic reconstructions acquired under low magnification with consecutive reacquisition of EM tomo box under high magnification and its re-examination was used exactly as described (Beznoussenko & Mironov, 2015). Briefly, an ultramicrotome (Leica EM UC7; Leica Microsystems, Vienna) was used to cut 200 nm serial semi-thick sections. Sections were collected on 1% Formvar films adhered to slot grids. Both sides of the grids were labelled with fiduciary 10 nm gold particles (PAG10, CMC, Utrecht, the Netherlands). Tilt-series were collected from the samples from ±65° with 1° increments at 200 kV in Tecnai 20 electron microscopes (FEI, Thermo Fisher Scientific, Eindhoven, the Netherlands). Tilt series were recorded at a magnification of 19,000x, using software supplied with the instrument. The nominal resolution in our tomograms was 4 nm, based upon section thickness, the number of tilts, tilt increments, and tilt angle range. The IMOD package and its newest viewer, 3DMOD 4.0.11, were used to construct individual tomograms and for the assignment of the outer leaflet of organelle membrane contours, CLEM was performed exactly as described (Beznoussenko *et al*, 2014).

### Stable EMD-EGFP PC-3 cell line generation

PC-3 cells were transfected with Emerin-pEGFP-C1 (Addgene plasmid #61993). Forty-eight hours post-transfection cells were cultured in medium containing G-418 (400 μg/ml, Life Technologies, Cat. 11811-031) for 1 week. Afterwards, single cells were plated onto 96-well plates. Colonies obtained from single cell expressing green fluorescence signal were selected and further propagated.

### Microscopy

Widefield fluorescent microscopy imaging was performed with Zeiss Axio Observer 7 equipped with Axiocam 506 monchromatic camera with 20x objective (Plan-Neofluar 20x/0.50). Z-stack images were acquired with confocal laser scanning microscope Leica TCS SP8 microscope (Leica Microsystems, Germany) with distance of 0.3-0.5 μm of focal plane using z-stack function. Oil immersion objectives were used: 20x oil immersion HC PL APO 0.75 NA and 63x oil immersion HC PL APO 1.4 NA objectives (Leica Microsystems, Germany). The microscope was equipped with software Leica Application Suite X (ver. 3.5.7.23225).

### Video-microscopy

Video-microscopy was performed on Leica TCS SP8 x microscope using incubation chambers and gas controllers from Life Imaging Services GmbH (Switzerland). PC-3 cells stably expressing Emerin-pEGFP-C1 plasmid (Addene, #61993) were left to attach on collagen-coated 35 mm glass-bottom dishes (MatTek Life Sciences) for 48h. Cells were imaged for 16-48h with 5 min. intervals. Oil immersion objectives were used: 20x oil immersion HC PL APO 0.75 NA and 63x oil immersion HC PL APO 1.4 NA objectives (Leica Microsystems, Germany). The microscope was equipped with software Leica Application Suite X (ver. 3.5.7.23225).

### siRNA transfection and drug treatement

Knock-down of selected targets was performed using siRNA (**Supplementary Table S3**) with the RNAiMax (Thermo Fisher Scientific #13778150) reagent following the manufactureŕs. Cells were cultured for 72h after transfection. Cells were treated by ICRF-193 (ChemCruz # CAS 21416-68-2) in concentration 5μM for 24h before analysis. CENP-E inhibitor (MCE inhibitors, # GSK-923295) was added in concentration 100nM for 8h and then wash out.

### Western blot

Proteins were obtained by cell lysis using RIPA buffer (Thermo Fisher Scientific #89901) containing Protease Inhibitor (Thermo Fisher Scientific #78438) inhibitors. Cleared protein lysates were separated on SDS-PAGE and transferred on nitrocellulose membranes. Membranes were incubated in 5 % skim milk in TBS-T as blocking reagent and incubated with primary antibodies (**Supplementary Table S2**) and appropriate secondary antibodies conjugated with fluorochromes. Protein bands were visualized on Odyssey CLx Imager.

### Gene knock-out generation

Lentivirus- CRISPR/Cas9-methodology mediated gene knock-outs were generated as described previously in (Wenta *et al*, 2022). Two independent, specific sequences targeting constitutive early exons of EMD were designed (EMD1 – TTGTACCGGCGCAGCAAGG targeted in exon 1, EMD2-TCTTCGAGTACGAGACCCAG targeted in exon 2). To prevent the off-target sites targeting, the specificity of gRNA sequences was validated using FASTA similarity search tool (EMBL-EBI) (more than three mismatches with any other site in the human genome (GRCh38.p13)). EMD1 and EMD2 gRNA sequences were inserted via BsmBI restriction enzyme into plentiCRISPRv2_puro vector (Addgene, 52961) and applied for lentivirus preparation. Lentiviral particles were generated by co-transfecting 5 μg of the specific plentiCRISPRv2, 3.75 μg psPax2, and 1.25 μg pVSV-G (second-generation lentivirus packaging system) into human embryonic kidney packaging cells GP2-293 (Clontech, Inc., Mountain View, CA, USA), using CalPhos™ Mammalian Transfection Kit (Clontech, Inc., Mountain View, CA, USA), according to the manufacturer’s protocol as described (Wenta *et al*, 2019). Positively transduced cells were selected with 1 μg/ml puromycin for at least 10 days and then analyzed by western blotting.

### Inverse fluorescence recovery after photobleaching (iFRAP)

Prostate cancer cell line PC-3 cells that expressed EMD-EGFP were left to attach on collagen-coated 35 mm glass-bottom dishes (MatTek Life Sciences) for 48h. The cells were imaged using an Olympus Spinning Disk CSU system based on an Olympus IX83 inverted microscope equipped with an Andor iXon Ultra camera. The images were acquired with a U PLAN S APO 60x/1.35 NA oil immersion objective using CellSens software (Olympus). NE and EMD-rich structures were photobleached using a 405nm laser at maximum power. The cells were imaged before and 7 minutes after the photobleaching with 5s timeframe. During the experiment the cells were maintained at 37C in humified atmosphere and 5% CO2 using an incubator system (Okolab).

### DAPI content quantification

For the analysis of the differential signal intensity in the nucleus and micronucleus, images were acquired, and nuclei and micronuclei were identified based on the DAPI and EMD channel. The DAPI signal was segmented using the Otsu filter (value=25). The mean signal intensity of region of interest, including nucleus and micronucleus was compared and the ratio between the two was calculated.

### Pauperization NE quantification

For the analysis of the NE pauperization, images were acquired, and cells with and without EMD-rich MN were identified. Based on the EMD channel and the signal was segmented using the Intermodes filter (value=25). The mean signal intensity of region of the whole NE was calculated and divided by the median cytoplasmic signal in cells without MN and cells with EMD-rich MN.

### Nuclear area measurement

The nuclei were identified as DAPI-positive areas, the signal was segmented using the Otsu filter. Their area and shape parameters were calculated with the identified and counted using structures using ImageJ’s Analyze Particles (https://imagej.net/imaging/particle-analysis).

### F-actin coherency

F-actin coherency was calculated from the projected confocal images using Orientation J plugin (http://bigwww.epfl.ch/demo/orientationj/) for ImageJ (Püspöki *et al*, 2016), where F-actin fibers coherency parameter was calculated in selected cells.

### Focal adhesion size analysis

In order to visualize focal adhesions, cells were stained with anti-paxillin antibody. Using the custom-built ImageJ macro (Nastały *et al*, 2020) the best z-plane was selected, the signal was segmented using Intermodes (value 25) and the size, numer and shape descriptors of the particles was measured. For each cell, an average area of focal adhesions was calculated.

### MTOC-nucleus distance quantification

Immunofluorescent staining using anti-pericentrin antibody was performed and nuclei were visualized using DAPI. After projecting Z-stacks, nuclei were registered to the reference nucleus using the turboreg ImageJ plugin. For each channel all the registered images were combined in a single stack. Then, the stack was segmented using threshold value for each image in the stack, using Yen (value 25) for pericentrin signal and Otsu (value 100) for nucleus. Finally, the custom-built ImageJ macro (Nastały *et al*, 2020) measured distance between the centrosome and the closest border of the nucleus for each cell. Value <0 means that centrosome was positioned above the nucleus.

### Nanostring nCounter Gene Expression Assay

Analysis of differentially expressed genes in relation to EMD status was prepared and performed as described (Nastały *et al*, 2021). Briefly, total mRNA was extracted from 69 samples prepared as formalin-fixed paraffin-embedded (FFPE) primary prostate tumor tissue cores (three 20 µm-thick, unstained FFPE sections per patient) using the Rneasy Mini Kit (Qiagen, Hilden, Germany) according to the manufacturer’s protocol. RNA integrity was assessed using the Agilent 2100 Bioanalyzer (Agilent Technologies, Santa Clara, CA, USA) with the Agilent RNA 6000 Pico Kit (Agilent Technologies). Extracted RNA (4 µL) was pre-amplified using the nCounter Low RNA Input Kit (NanoString Technologies, Seattle, WA, USA) with the dedicated Primer Pool covering the sequences of 730 immune-related genes included in the nCounter Cancer Progression Profiling Panel (NanoString Technologies). Pre-amplified samples were analyzed using the NanoString nCounter Analysis System (NanoString Technologies) according to the manufacturer’s procedures for hybridization, detection, and scanning.

Background correction and data normalization were performed using nSolver 4.0 software (NanoString Technologies). Background level was estimated by thresholding over the mean plus 2 standard deviations of negative control counts, and data were normalized according to the global mean of the counts of positive control probes included in the assay and 4 most stably expressed housekeeping genes – CNOT4, HDAC3, DDX50 and CC2D1B. Sixty-six primary prostate tumors were available for EMD imaging analysis (uninformative or technically damaged samples were excluded from the analyses) and their transcriptomic profile was further correlated to the EMD status. Data normalization, housekeeping genes, stratification (EMD cutoffs were <25% percentile value: <0.07538 EMD-rich MN/nucleus; median: 0.1280 EMD-rich MN/nucleus; >75% percentile value: 0.1861). Emerin negative PCa (pauperized EMD) was defined as >85% tumor cells negative or low staining intensity (+1) for EMD staining.

### Spheroid culture

Single PC-3 cells were resuspended in growth factor reduced solubilized basement membrane (Matrigel®, Corning) mixed with Collagen I (Corning® Collagen I, High Concentration, Rat Tail) to obtain concentration of 5 mg/mL and plated into drops in 8-well glass-bottom ibidi chambers (Ibidi) at 8000 cells/100 μL. They were grown for 7 days in RPMI 1640 (Thermo Fisher Scientific #11875093) supplemented with 10% FBS (Thermo Fisher Scientific #10270-106), medium was changed every 2 days The spheroids were fixed with 4%PFA and stained overnight with primary antibodies diluted in 0.5% Triton X-100/5%FBS, followed by the secondary antibody incubation in the same buffer. The single spheroids were then imaged with confocal microscopy. The invading cell is determined as cell that had the nucleus positioned out the main cell mass. The properties of spheroids including area, circularity and solidity were quantified using ImageJ.

### RNA sequencing

The total RNA from sferoides was extracted using PureLink™ RNA Mini Kit (ThermoFisher Sci.) according to the manufacturer’s instructions following Phenol/Chloroform purification (ThermoFisher Sci.) RNA purity and concentration testing was performed using the Nanodrop2000 (Life Technologies). Agilent RNA ScreenTape System (Agilent) were used to RNA integrity analysis. RNA sequencing libraries were generated according to the manufacturer’s instructions for the TruSeq totalRNA with RiboZero Human/Mouse/Rat Gold (Illumina, San Diego, CA, United States). Sequencing was then performed on the NovaSeq6000 (Illumina, San Diego, CA, United States) platform.

### Time-lapse microscopy and migration analysis

Prostate cancer PC-3 cells expressing Emerin-pEGFP-C1 plasmid (Addene, #61993) were imaged using Operetta CLS microscope in confocal mode with a 40x water Immersion lens (NA 1.1). Images were acquired every 7 minutes for 16 h. The cells were then manually tracked using the ImageJ TrackMate plugin (Ershov *et al*, 2022).

### Transwell assay

Transwells (pore size of 8.0 μm, Corning) were coated with collagen I (Corning® Collagen I, High Concentration, Rat Tail) for migration assays. Briefly, 1 × 10^5^ cells were resuspended in 500 μL of serum-free medium and added to the top compartment of a transwell chamber. Five-hundred microliters of medium supplemented with 10% FBS were added to a 12-well plate. The transwell was placed on the plate for 24 h. The number of EMD-rich MN/cells that migrated through the pore were quantified by imaging using at least 4 randomly chosen fields/experiment.

### Collagen-invasion assay

PC-3 cells were seeded at optimal density (30,000 cells/well) and allowed to adhere before removal of media. 40 μL Collagen I (Corning® Collagen I, High Concentration, Rat Tail) was overlaid at 5 mg/mL. Fresh RPMI 1640 (Thermo Fisher Scientific #11875093) supplemented with 10% FBS (Thermo Fisher Scientific #10270-106) was added, and cell invasion was assayed after incubation for 48h by fixing cells with 4%PFA followed by IF staining. The number of EMD-rich MN/cells that invaded the collagen were quantified by imaging using at least 5 randomly chosen fields/experiment and compared to the cells that did not invaded the gel.

### TCGA and cBio portal data

The Cancer Genome Atlas (TCGA) prostate adenocarcinoma (PRAD) clinical characteristics and RNA-seq data (RNASeqV2, RSEM normalized) covering normalized counts of sequences aligning to 20,531 genes were obtained for 497 PRAD patients from cBioPortal for Cancer Genomics (E *et al*, 2012) (Firehose Legacy dataset, data access: 26^th^ July 2022). The methods of biospecimen procurement, RNA isolation and RNA sequencing were previously described by TCGA Research Network (The molecular taxonomy of primary prostate cancer, 2015). EMD mutation data were obtained from the cBio Porta for Cancer Genomics (https://www.cbioportal.org/), where 1,607 of prostate adenocarcinoma samples profiled for mutations the *EMD* gene were available (data access: 6^th^ of September 2023).

### Bioinformatic analyses

*RNA expression in PC-3 cell line.* See supplementary Information: Material and Methods.

*PCa tumors*. In nCounter gene expression dataset, differentially expressed genes (DEGs) were identified based on p < 0.05 (Mann-Whitney-Wilcoxon test) and median-based log2FC values for pauperized vs normal emerin comparison were reported. DEGs were associated with GO BP (Gene Ontology Biological Process) terms using Functional Annotation Tool by DAVID Bioinformatics Resources 6.8 (Huang *et al*, 2007a, 2007b) and the interaction network for their protein products was visualized using STRING v11(Szklarczyk *et al*, 2019).In TCGA PRAD cohort, EMD-pauperized transcriptomic score was calculated as mean of log2 RSEM-normalized counts of 9 genes (*CXCR4*, *APOE*, *SPARC*, *VIM*, *GSN*, *ANXA2P2*, *SFRP1*, *COL18A1*, *FN1*) in each sample. EMD-pauperization status was categorized according to upper quartile of the score.

### Statistical analysis and reproducibility

All data are presented as scatter plots or box plots expressed as mean ± SD, unless otherwise indicated. The number of experiments as well as the number of samples analyzed is specified for each experiment and additional methods used to calculate these are described in the figure legends are reported in the figure legends. Statistical significance was calculated, whenever we compared two distinct distributions, using a parametric two-tails unpaired student’s t-test with unpaired t-test, paired t-test or one-way ANOVA with Dunnett’s, corrections or non-parametric two-tailed Mann–Whitney U t-test as indicated. Kruskal–Wallis/Dunn’s test was used for one-way data with more than two groups. Paired analysis was performed using Wilcoxon matched-pairs signed rank test. Contingency analysis was performed using Fisher’s Exact test. Associations between frequency of EMD-rich MN and time-to-biochemical recurrence were evaluated using log-rank test and presented as Kaplan–Meier plots. *p < 0.05, **p < 0.01, ***p < 0.001, ****p < 0.0001. Statistical analysis of quantitative data was carried out in GraphPad Prism v. 9.0.0. or in R (4.2.2. version) using R statistical packages. Graphic representation was prepared with Biorender.

## List of abbreviations

BR: biochemical recurrence
BRCA2: BRCA2, DNA repair associated
CLEM: Correlative light electron microscopy
DFS: disease-free survival
EMD-NE level MN: Emerin-Nuclear envelope level micronucleus
EMD-rich MN: Emerin-rich micronucleus
EMD: Emerin
EGFP: Enhanced green fluorescence protein
ER: Endoplasmic reticulum
FA: focal adhesion
iFRAP: inverse fluorescence recovery after photobleaching
KD: knock-down
KO: knock-out
LBR: Lamin B receptor
LAP2a: Lamina-associated polypetide
LEM-domain: LAP2-Emerin-MAN1
MN: Micronucleus
MTOC: microtubule-organizing center
NE: Nuclear envelope
PCa: prostate cancer
PSA: prostate-specific antigen
STED: Super resolution Stimulated Emission Depletion
TGCA: The Cancer Genome Atlas
TMA: Tissue Microarray

## Declarations

### Ethics approval and consent to participate

The study has been approved by the local Ethics Committee (i.e., Independent Bioethics Committee for Scientific Research at Medical University of Gdańsk, no. NKBBN/286/2018). Written informed consent was obtained from all the subjects involved in the study.

### Consent for publication

All authors have agreed with publishing this manuscript.

### Availability of data and materials

The datasets used and/or analyzed in this article were included within the article and the additional files. All data are available in the main text or the supplementary materials. Please contact the corresponding author for data requests.

### Authors’ contributions

Conceptualization and study design: PN

Results interpretation: PN, PM

Data analyses: PN, MP, KK, TW

Methodology: PN, MP, KK, NBK, JR, GVB, MR, TW, AM, ZL, SB, LB, AŻ, RB

Clinical samples design: NBK

Clinical sample acquisition: JR, JSz, KM, MS, MM, NBK

Data curation: PN, JR, KM, NBK

Investigation: PN, KK, MP

Visualization: PN, KK, MP

Funding acquisition: PN

Project administration: PN

Supervision: PN

Writing – original draft: PN

Writing – review & editing: PN, PM, MP, NBK, GDS

## Acknowledgments

The authors would like to thank all the patients that participated in this study. The authors would like to also thank Dario Parazzoli for the access to IFOM Imaging Facility. We would like to express gratitude to Kamil Myszczyński for RNAseq dataset preparation.

## Extended Data Figures

**Extended Data Figure 1.**
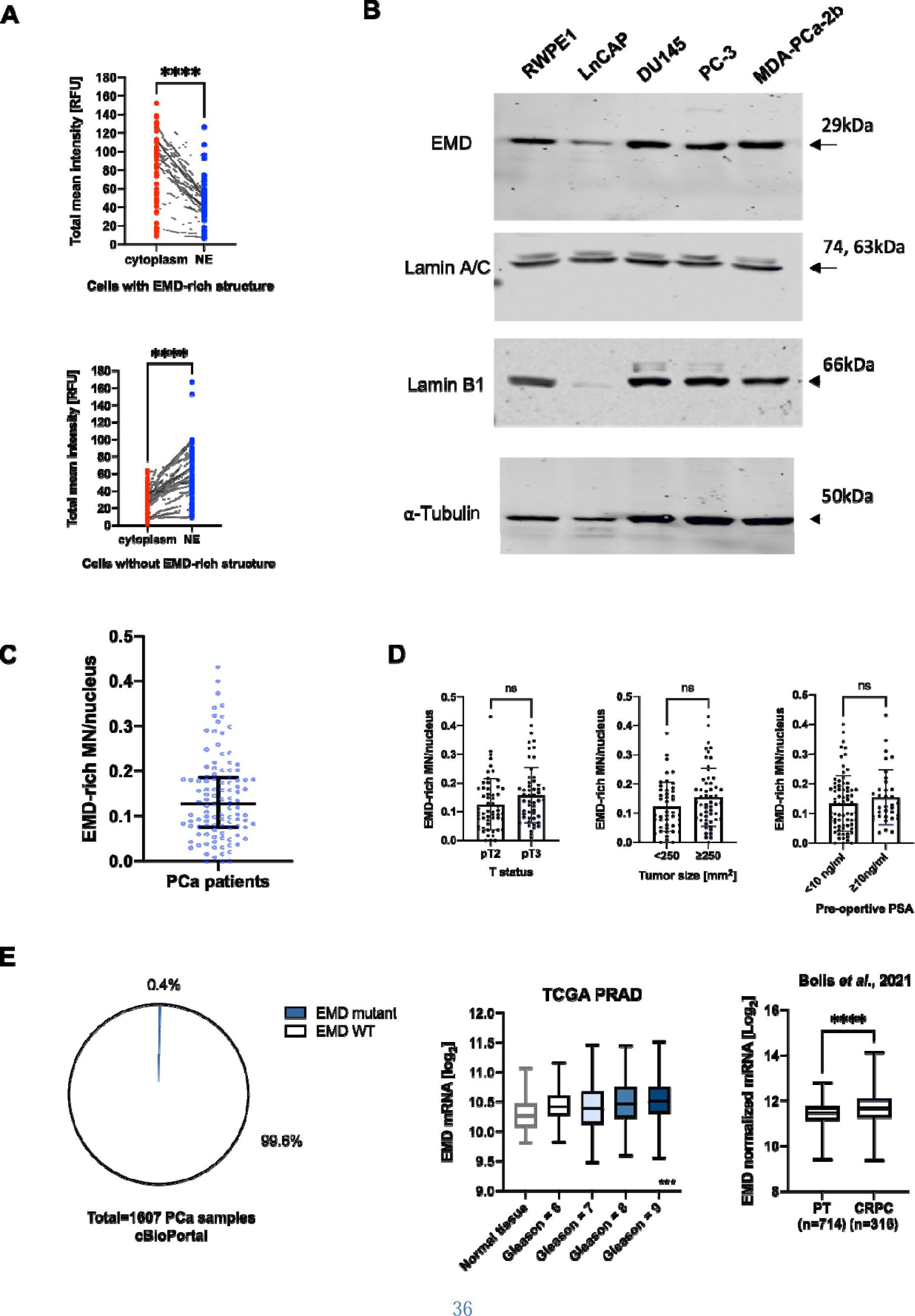
**a** Total mean EMD intensity paired measurement in NE comparing to cytoplasm cells with EMD-rich MN (n=51 cells), and cells without EMD-rich MN (n-63 cells). **b** Western blot performed in various prostate cell lines for EMD, Lamin A/C and Lamin B1. **c** EMD-rich structures distribution among PCa patients (n=107) **d** Clinical associations between EMD-rich MN and T status (n=107 patients) tumor size (n=100 patients), and pre-operative PSA levels (n=119 patients). **e** Prevalence of EMD mutations in prostate adenocarcinoma according to cBio Portal for Cancer Genomics; association of EMD mRNA expression to Gleason score according to The Cancer Genome Atlas; comparison between EMD mRNA expression in primary prostate cancer tumors (PT) and castration-resistant prostate cancer (CRPC) according to Bolis *et al.,* 2021. Paired plot: Wilcoxon matched-pair signed rank test; Box-plot; Whiskers indicate min to max values, within the box the first quartile, median, third quartile are represented. For multiple comparison Kruskal-Wallis test was used. Scatter-plot with bar (mean with SD) and bar-plots Mann-Whitney U test. Error bars indicate SD.*P < 0.05; **P < 0.01; ***P < 0.001; ****P < 0.0001; ns, not significant. Source data is provided as Source data file.

**Extended Data Figure 2.**
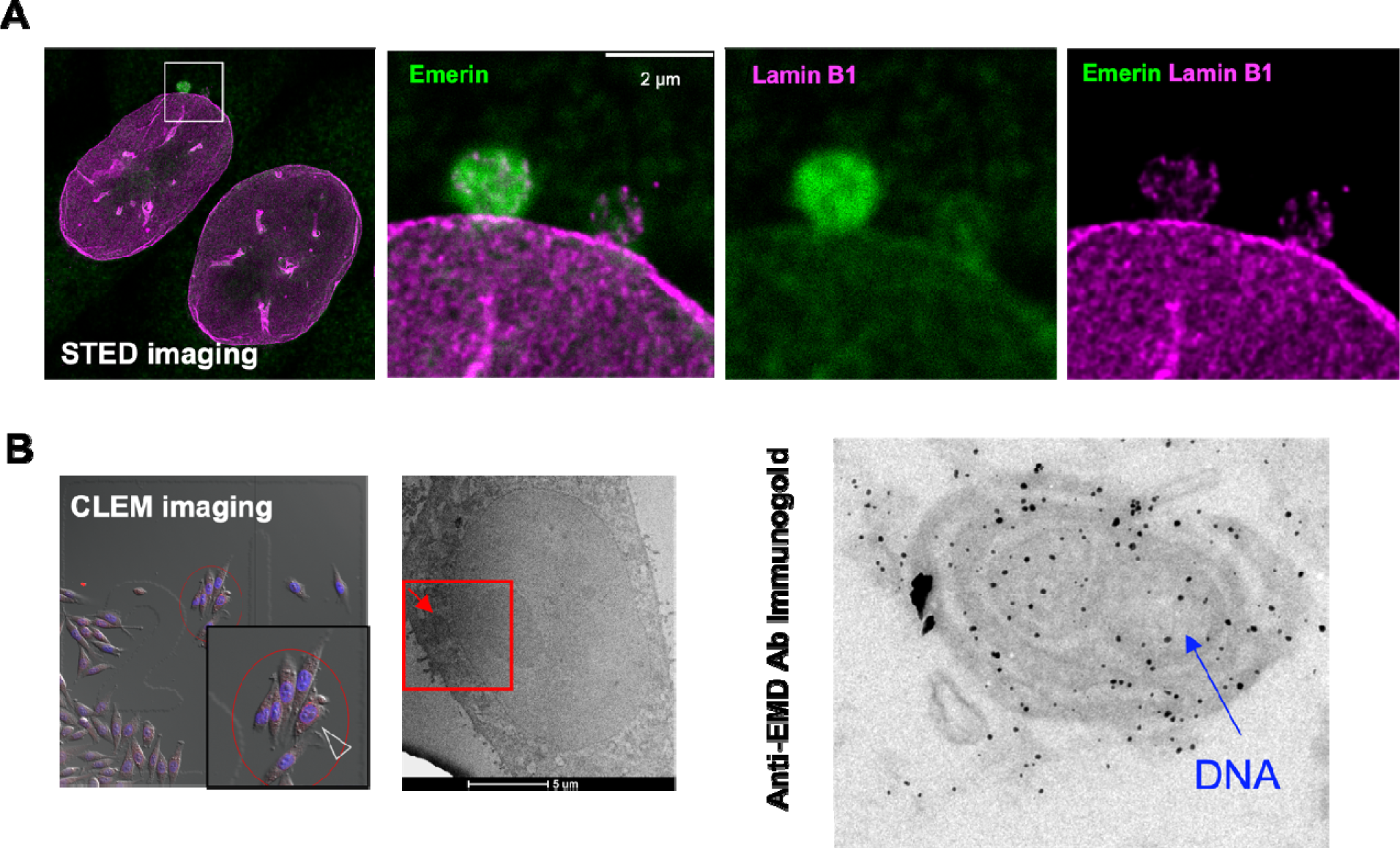
**a** Super Resolution Stimulated Emission Depletion (STED) microscopy of EMD-rich MN stained for EMD and Lamin B1, EMD is shown in green, Lamin B1 in magenta. **b** Brightfield image and electron microscopy image of a cell in Correlative light electron microscopy (CLEM) experiment of EMD-rich MN. Left panel, immunogold labeling of EMD-rich MN.EXt

**Extended Data Figure 3.**
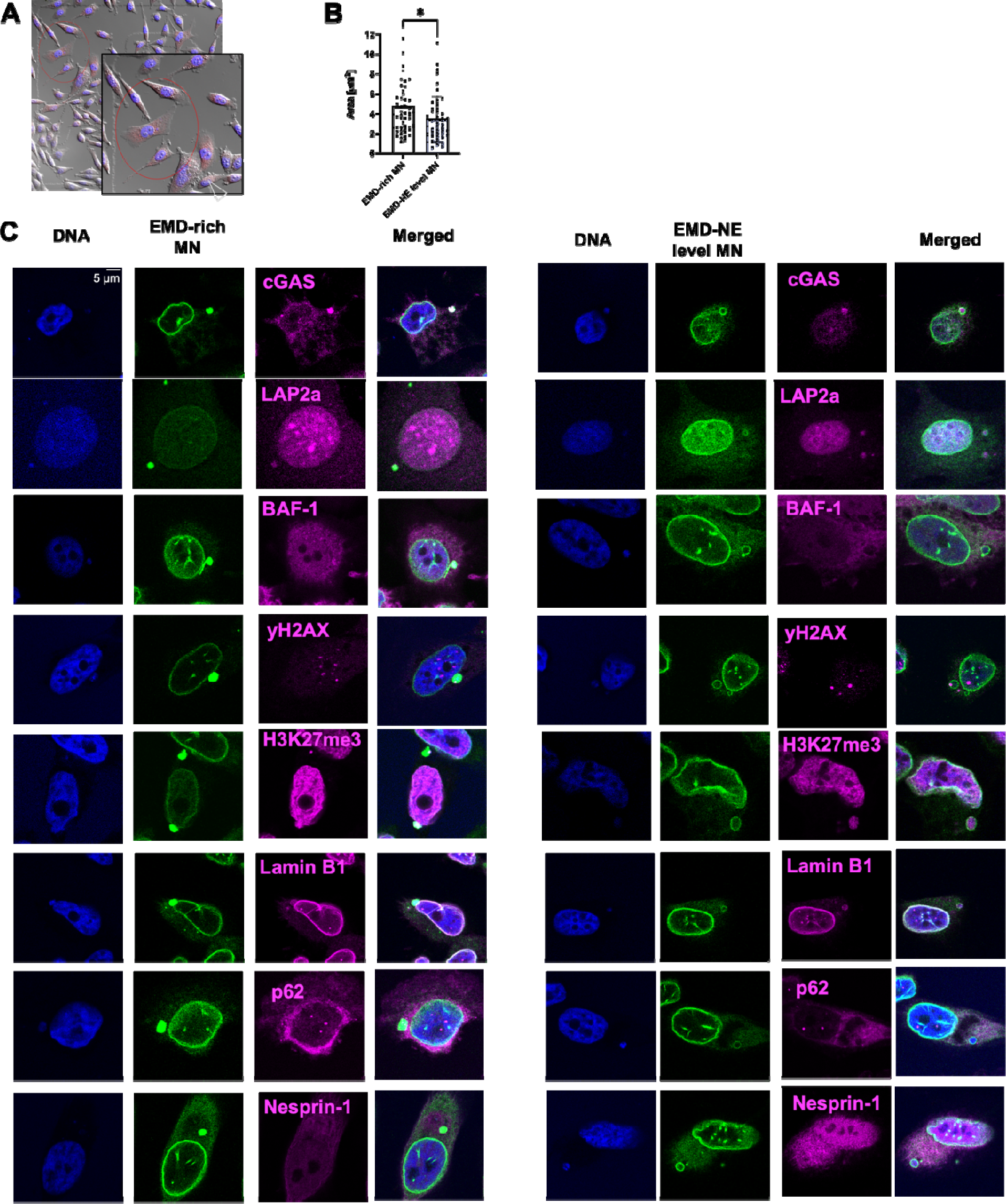
**a** Brightfield image of a cell with EMD-rich MN and EMD-NE-level MN that was further used in correlative light electron microscopy (CLEM). **b** Area quantification of EMD-rich MN (n=51 cells) and EMD-NE-level MN (n=50 cells). **c** Representative micrographs of staining for cGAS, LAP2a, BAF-1, yX2AX, H3K27me2, Lamin B1, p62, and Nesprin-1 in EMD-rich MN and EMD-NE-level MN, EMD is presented in green, other proteins in magenta. Scatter-plot with bar (mean with SD) and bar-plots Mann-Whitney U test. Error bars indicate SD.*P < 0.05; **P < 0.01; ***P < 0.001; ****P < 0.0001; ns, not significant. Source data is provided as Source data file.

**Extended Data Figure 4.**
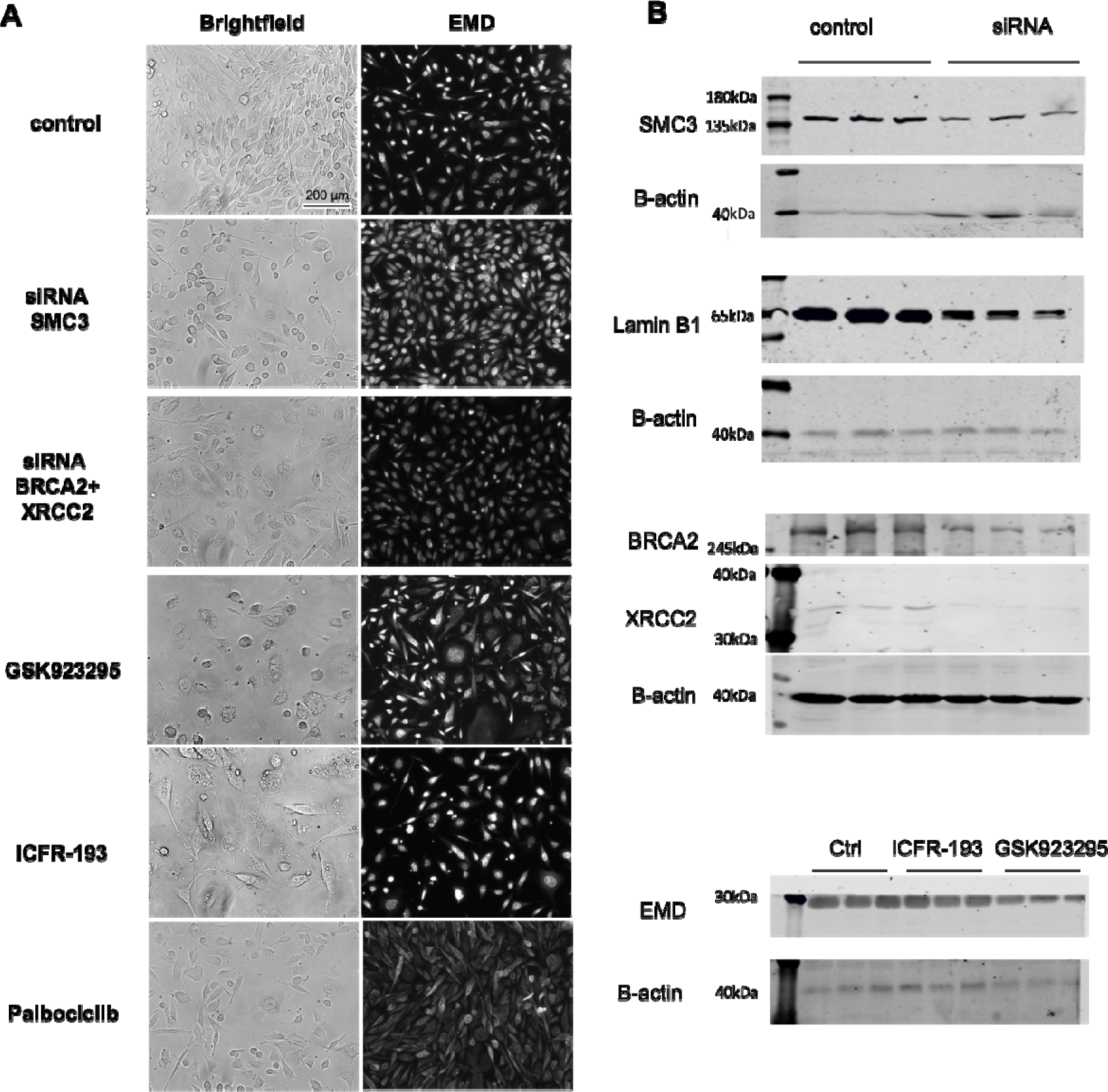
**a** Representative images of brightfield and EMD in control PC-3 cells and the treated ones. **b** Western blot analysis of EMD in PC-3 cells under various knock-down conditions. Source data is provided as Source data file.

**Extended Data Figure 5.**
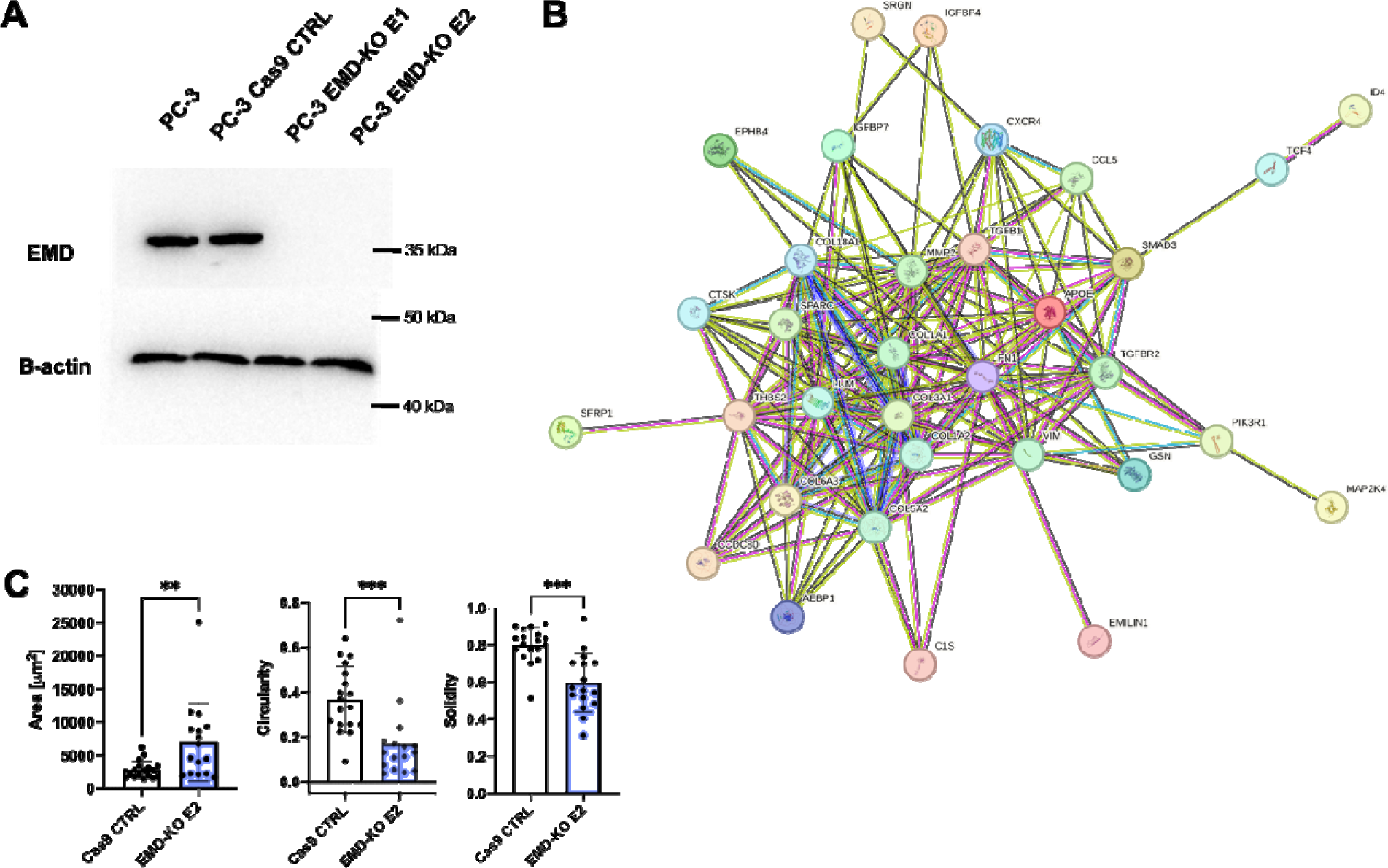
**a** The interaction network of protein products og differentially expressed genes in PCa tumors with EMD pauperized phenotype visualized using STRING v11. **b** Western blot analysis of EMD in PC-3 control and EMD-KO cells. **c** Comparison of properties including area, circularity, solidity of PC-3 control (n=18) and EMD-KO (n=16) spheroids. Scatter-plot with bar (mean with SD) and bar-plots Mann-Whitney U test. Error bars indicate SD.*P < 0.05; **P < 0.01; ***P < 0.001; ****P < 0.0001; ns, not significant. Source data is provided as Source data file.

## Supplementary Information

Supplementary Material and Methods

Supplementary Tables S1 to S5

Supplementary Movies 1-6

## References

Agustinus AS, Al-Rawi D, Dameracharla B, Raviram R, Jones BSCL, Stransky S, Scipioni L, Luebeck J, Di Bona M, Norkunaite D, et al (2023) Epigenetic dysregulation from chromosomal transit in micronuclei. Nature 619: 176–183

Bankhead P, Loughrey MB, Fernández JA, Dombrowski Y, McArt DG, Dunne PD, McQuaid S, Gray RT, Murray LJ, Coleman HG, et al (2017) QuPath: Open source software for digital pathology image analysis. Sci Rep 7: 16878

Bednarz-Knoll N, Eltze E, Semjonow A & Brandt B (2019) BRCAness in prostate cancer. Oncotarget 10: 2421

Berk JM, Simon DN, Jenkins-Houk CR, Westerbeck JW, Grønning-Wang LM, Carlson CR & Wilson KL (2014) The molecular basis of emerin–emerin and emerin–BAF interactions. Journal of Cell Science 127: 3956–3969

Beznoussenko GV & Mironov AA (2015) Correlative video-light-electron microscopy of mobile organelles. Methods Mol Biol 1270: 321–346

Beznoussenko GV, Parashuraman S, Rizzo R, Polishchuk R, Martella O, Di Giandomenico D, Fusella A, Spaar A, Sallese M, Capestrano MG, et al (2014) Transport of soluble proteins through the Golgi occurs by diffusion via continuities across cisternae. eLife 3: e02009

Bolis M, Bossi D, Vallerga A, Ceserani V, Cavalli M, Impellizzieri D, Di Rito L, Zoni E, Mosole S, Elia AR, et al (2021) Dynamic prostate cancer transcriptome analysis delineates the trajectory to disease progression. Nat Commun 12: 7033

Brachner A & Foisner R (2014) Lamina-associated polypeptide (LAP)2α and other LEM proteins in cancer biology. Adv Exp Med Biol 773: 143–163

Buchwalter A, Schulte R, Tsai H, Capitanio J & Hetzer M (2019) Selective clearance of the inner nuclear membrane protein emerin by vesicular transport during ER stress. eLife 8: e49796

Chang W, Folker ES, Worman HJ & Gundersen GG (2013) Emerin organizes actin flow for nuclear movement and centrosome orientation in migrating fibroblasts. Mol Biol Cell 24: 3869–3880

Cimini D, Moree B, Canman JC & Salmon ED (2003) Merotelic kinetochore orientation occurs frequently during early mitosis in mammalian tissue cells and error correction is achieved by two different mechanisms. J Cell Sci 116: 4213–4225

Crasta K, Ganem NJ, Dagher R, Lantermann AB, Ivanova EV, Pan Y, Nezi L, Protopopov A, Chowdhury D & Pellman D (2012) DNA breaks and chromosome pulverization from errors in mitosis. Nature 482: 53–58

Crozier L, Foy R, Mouery BL, Whitaker RH, Corno A, Spanos C, Ly T, Cook JG & Saurin AT (2022) CDK4/6 inhibitors induce replication stress to cause longLJterm cell cycle withdrawal. The EMBO Journal 41

Dechat T, Gajewski A, Korbei B, Gerlich D, Daigle N, Haraguchi T, Furukawa K, Ellenberg J & Foisner R (2004) LAP2α and BAF transiently localize to telomeres and specific regions on chromatin during nuclear assembly. Journal of Cell Science 117: 6117–6128

Dhital B, Santasusagna S, Kirthika P, Xu M, Li P, Carceles-Cordon M, Soni RK, Li Z, Hendrickson RC, Schiewer MJ, et al (2023) Harnessing transcriptionally driven chromosomal instability adaptation to target therapy-refractory lethal prostate cancer. Cell Reports Medicine 4: 100937

E C, J G, U D, Be G, So S, Ba A, A J, Cj B, Ml H, E L, et al (2012) The cBio cancer genomics portal: an open platform for exploring multidimensional cancer genomics data. Cancer discovery 2

Ershov D, Phan M-S, Pylvänäinen JW, Rigaud SU, Le Blanc L, Charles-Orszag A, Conway JRW, Laine RF, Roy NH, Bonazzi D, et al (2022) TrackMate 7: integrating state-of-the-art segmentation algorithms into tracking pipelines. Nat Methods 19: 829–832

Fenech M, Kirsch-Volders M, Natarajan AT, Surralles J, Crott JW, Parry J, Norppa H, Eastmond DA, Tucker JD & Thomas P (2011) Molecular mechanisms of micronucleus, nucleoplasmic bridge and nuclear bud formation in mammalian and human cells. Mutagenesis 26: 125–132

Ferrandiz N, Downie L, Starling GP & Royle SJ (2022) Endomembranes promote chromosome missegregation by ensheathing misaligned chromosomes. J Cell Biol 221: e202203021

Flynn PJ, Koch PD & Mitchison TJ (2021) Chromatin bridges, not micronuclei, activate cGAS after drug-induced mitotic errors in human cells. Proc Natl Acad Sci USA 118: e2103585118

Hale CM, Shrestha AL, Khatau SB, Stewart-Hutchinson PJ, Hernandez L, Stewart CL, Hodzic D & Wirtz D (2008) Dysfunctional Connections Between the Nucleus and the Actin and Microtubule Networks in Laminopathic Models. Biophysical Journal 95: 5462–5475

Hatch EM, Fischer AH, Deerinck TJ & Hetzer MW (2013) Catastrophic nuclear envelope collapse in cancer cell micronuclei. Cell 154: 47–60

Hell SW (2007) Far-Field Optical Nanoscopy. Science 316: 1153–1158

Huang DW, Sherman BT, Tan Q, Collins JR, Alvord WG, Roayaei J, Stephens R, Baseler MW, Lane HC & Lempicki RA (2007a) The DAVID Gene Functional Classification Tool: a novel biological module-centric algorithm to functionally analyze large gene lists. Genome Biology 8: R183

Huang DW, Sherman BT, Tan Q, Kir J, Liu D, Bryant D, Guo Y, Stephens R, Baseler MW, Lane HC, et al (2007b) DAVID Bioinformatics Resources: expanded annotation database and novel algorithms to better extract biology from large gene lists. Nucleic Acids Research 35: W169–W175

Jiang H, Kong N, Liu Z, West SC & Chan YW Human Endonuclease ANKLE1 Localizes at the Midbody and Processes Chromatin Bridges to Prevent DNA Damage and cGAS-STING Activation. Advanced Science n/a: 2204388

Kim D-H & Wirtz D (2013) Focal adhesion size uniquely predicts cell migration. FASEB J 27: 1351–1361

Kneissig M, Keuper K, de Pagter MS, van Roosmalen MJ, Martin J, Otto H, Passerini V, Campos Sparr A, Renkens I, Kropveld F, et al (2019) Micronuclei-based model system reveals functional consequences of chromothripsis in human cells. eLife 8: e50292

Kwon M, Leibowitz ML & Lee J-H (2020) Small but mighty: the causes and consequences of micronucleus rupture. Exp Mol Med 52: 1777–1786

Lammerding J, Hsiao J, Schulze PC, Kozlov S, Stewart CL & Lee RT (2005) Abnormal nuclear shape and impaired mechanotransduction in emerin-deficient cells. J Cell Biol 170: 781–791

Lang SH, Hyde C, Reid IN, Hitchcock IS, Hart CA, Bryden AAG, Villette J-M, Stower MJ & Maitland NJ (2002) Enhanced expression of vimentin in motile prostate cell lines and in poorly differentiated and metastatic prostate carcinoma. Prostate 52: 253–263

Lavenus SB, Vosatka KW, Caruso AP, Ullo MF, Khan A & Logue JS (2022) Emerin regulation of nuclear stiffness is required for fast amoeboid migration in confined environments. J Cell Sci: jcs.259493

Leylek TR, Jeusset LM, Lichtensztejn Z & McManus KJ (2020) Reduced Expression of Genes Regulating Cohesion Induces Chromosome Instability that May Promote Cancer and Impact Patient Outcomes. Sci Rep 10: 592

Liddane AG & Holaska JM (2021) The Role of Emerin in Cancer Progression and Metastasis. Int J Mol Sci 22: 11289

Liddane AG, McNamara CA, Campbell MC, Mercier I & Holaska JM (2021) Defects in Emerin-Nucleoskeleton Binding Disrupt Nuclear Structure and Promote Breast Cancer Cell Motility and Metastasis. Mol Cancer Res 19: 1196–1207

Liu S, Kwon M, Mannino M, Yang N, Renda F, Khodjakov A & Pellman D (2018) Nuclear envelope assembly defects link mitotic errors to chromothripsis. Nature 561: 551–555

López-Moncada F, Torres MJ, Lavanderos B, Cerda O, Castellón EA & Contreras HR (2022) SPARC Induces E-Cadherin Repression and Enhances Cell Migration through Integrin αvβ3 and the Transcription Factor ZEB1 in Prostate Cancer Cells. Int J Mol Sci 23: 5874

Lu S, Lee J, Revelo M, Wang X, Lu S & Dong Z (2007) Smad3 is overexpressed in advanced human prostate cancer and necessary for progressive growth of prostate cancer cells in nude mice. Clin Cancer Res 13: 5692–5702

Maass KK, Rosing F, Ronchi P, Willmund KV, Devens F, Hergt M, Herrmann H, Lichter P & Ernst A (2018) Altered nuclear envelope structure and proteasome function of micronuclei. Experimental Cell Research 371: 353–363

Maciejowski J, Li Y, Bosco N, Campbell PJ & de Lange T (2015) Chromothripsis and Kataegis Induced by Telomere Crisis. Cell 163: 1641–1654

Mackenzie KJ, Carroll P, Martin C-A, Murina O, Fluteau A, Simpson D, Olova N, Sutcliffe H, Rainger J, Robertson A, et al (2017) cGAS surveillance of micronuclei links genome instability to innate immunity. Nature 548: 461

Miller ET, You S, Cadaneanu RM, Kim M, Yoon J, Liu ST, Li X, Kwan L, Hodge J, Quist MJ, et al (2020) Chromosomal instability in untreated primary prostate cancer as an indicator of metastatic potential. BMC Cancer 20: 398

Molinié V, Fromont G, Sibony M, Vieillefond A, Vassiliu V, Cochand-Priollet B, Hervé JM, Lebret T & Baglin AC (2004) Diagnostic utility of a p63/α-methyl-CoA-racemase (p504s) cocktail in atypical foci in the prostate. Mod Pathol 17: 1180–1190

Nagano A, Koga R, Ogawa M, Kurano Y, Kawada J, Okada R, Hayashi YK, Tsukahara T & Arahata K (1996) Emerin deficiency at the nuclear membrane in patients with Emery-Dreifuss muscular dystrophy. Nat Genet 12: 254–259

Nastały P, Purushothaman D, Marchesi S, Poli A, Lendenmann T, Kidiyoor GR, Beznoussenko GV, Lavore S, Romano OM, Poulikakos D, et al (2020) Role of the nuclear membrane protein Emerin in front-rear polarity of the nucleus. Nature Communications 11: 2122

Nastały P, Smentoch J, Popęda M, Martini E, Maiuri P, Żaczek AJ, Sowa M, Matuszewski M, Szade J, Kalinowski L, et al (2021) Low Tumor-to-Stroma Ratio Reflects Protective Role of Stroma against Prostate Cancer Progression. J Pers Med 11: 1088

Procter DJ, Furey C, Garza-Gongora AG, Kosak ST & Walsh D (2020) Cytoplasmic control of intranuclear polarity by human cytomegalovirus. Nature 587: 109–114

Püspöki Z, Storath M, Sage D & Unser M (2016) Transforms and Operators for Directional Bioimage Analysis: A Survey. Adv Anat Embryol Cell Biol 219: 69–93

Reis-Sobreiro M, Chen J-F, Novitskaya T, You S, Morley S, Steadman K, Gill NK, Eskaros A, Rotinen M, Chu C-Y, et al (2018) Emerin Deregulation Links Nuclear Shape Instability to Metastatic Potential. Cancer Res 78: 6086–6097

Rowat AC, Lammerding J & Ipsen JH (2006) Mechanical properties of the cell nucleus and the effect of emerin deficiency. Biophys J 91: 4649–4664

Schindelin J, Arganda-Carreras I, Frise E, Kaynig V, Longair M, Pietzsch T, Preibisch S, Rueden C, Saalfeld S, Schmid B, et al (2012) Fiji: an open-source platform for biological-image analysis. Nat Methods 9: 676–682

Schmidt A, Kaakinen M, Wenta T & Manninen A (2022) Loss of α6β4 Integrin-Mediated Hemidesmosomes Promotes Prostate Epithelial Cell Migration by Stimulating Focal Adhesion Dynamics. Frontiers in Cell and Developmental Biology 10

Schneider CA, Rasband WS & Eliceiri KW (2012) NIH Image to ImageJ: 25 years of image analysis. Nat Methods 9: 671–675

Stich HF & Rosin MP (1984) Micronuclei in exfoliated human cells as a tool for studies in cancer risk and cancer intervention. Cancer Lett 22: 241–253

Stopsack KH, Whittaker CA, Gerke TA, Loda M, Kantoff PW, Mucci LA & Amon A (2019) Aneuploidy drives lethal progression in prostate cancer. Proceedings of the National Academy of Sciences 116: 11390–11395

Sung H, Ferlay J, Siegel RL, Laversanne M, Soerjomataram I, Jemal A & Bray F (2021) Global Cancer Statistics 2020: GLOBOCAN Estimates of Incidence and Mortality Worldwide for 36 Cancers in 185 Countries. CA: A Cancer Journal for Clinicians 71: 209–249

Szklarczyk D, Gable AL, Lyon D, Junge A, Wyder S, Huerta-Cepas J, Simonovic M, Doncheva NT, Morris JH, Bork P, et al (2019) STRING v11: protein–protein association networks with increased coverage, supporting functional discovery in genome-wide experimental datasets. Nucleic Acids Research 47: D607

The molecular taxonomy of primary prostate cancer (2015) Cell 163: 1011–1025

Thompson SL & Compton DA (2011) Chromosome missegregation in human cells arises through specific types of kinetochore-microtubule attachment errors. Proc Natl Acad Sci U S A 108: 17974–17978

Timms K, Cuzick J, Neff C, Reid J, Solimeno C, Sangale Z, Pruss D, Gutin A, Lanchbury J & Stone S (2016) The molecular landscape of genome instability in prostate cancer (PC). Annals of Oncology 27: vi35

Trudel D, Fradet Y, Meyer F, Harel F & Têtu B (2003) Significance of MMP-2 Expression in Prostate Cancer: an Immunohistochemical Study. Cancer Research 63: 8511–8515

Umbreit NT, Zhang C-Z, Lynch LD, Blaine LJ, Cheng AM, Tourdot R, Sun L, Almubarak HF, Judge K, Mitchell TJ, et al (2020) Mechanisms generating cancer genome complexity from a single cell division error. Science 368: eaba0712

Vietri M, Schultz S, Bellanger A, Jones C, Petersen L, Raiborg C, Skarpen E, Pedurupillay C, Kjos I, Kip E, et al (2020) Unrestrained ESCRT-III drives micronuclear catastrophe and chromosome fragmentation. Nature Cell Biology 22: 1–12

Wenta T, Rychlowski M, Jarzab M & Lipinska B (2019) HtrA4 Protease Promotes Chemotherapeutic-Dependent Cancer Cell Death. Cells 8: 1112

Wenta T, Schmidt A, Zhang Q, Devarajan R, Singh P, Yang X, Ahtikoski A, Vaarala M, Wei G-H & Manninen A (2022) Disassembly of α6β4-mediated hemidesmosomal adhesions promotes tumorigenesis in PTEN-negative prostate cancer by targeting plectin to focal adhesions. Oncogene 41: 3804–3820

