## Supplementary Information for "Pauperization of Emerin from nuclear envelope during chromatin bridge resolution drives prostate cancer cell migration and invasiveness"

Supplementary Material and Methods

Supplementary Tables S1 to S5

Caption for Supplementary Movies 1-6

**Supplementary Material and Methods**

*RNA expression in PC-3 cell line.* Data quality was checked using FastQC 0.11.9 [1] and MultiQC 1.11 [2] reporting tools. After quality control, raw reads were trimmed using Trimmomatic 0.39 [3] and then aligned to the human genome GRCh38.p10 using STAR 2.7.10 [4]. Mapped reads were counted across genomic features using featureCounts 2.0.3 [5]. Read counts were normalized and then subjected to differential analysis using DESeq2 1.360.0 [6]. The absolute value of log2 fold change ≥ (0.5) and adjusted p-value < 0.05 were used as a criteria to identify differentially expressed genes.

**References**

1. Andrews S, Krueger F, Seconds-Pichon A, Biggins F, Wingett S. FastQC. A quality control tool for high throughput sequence data. Babraham Bioinf. 2015;1:1.
2.Ewels P, Magnusson M, Lundin S, Kaller M. MultiQC: Summarize analysis results for multiple tools and samples in a single report. Bioinformatics. 2016;32:19.

3. Bolger AM, Lohse M, Usadel B. Trimmomatic: a flexible trimmer for illumina sequence data. Bioinformatics. 2014;30:2114-2120.
4. Dobin A, Davis CA, Schlesinger F, et al. STAR: ultrafast universal RNA-seq aligner. Bioinformatics. 2013;29:15-21.

5. Liao Y, Smyth GK, Shi W. featureCounts: an efficient general purpose program for assigning sequence reads to genomic features. Bioinformatics. 2014 Apr 1;30(7):923-30. doi:10.1093/bioinformatics/btt656. Epub 2013 Nov 13. PMID: 24227677.
6. Love MI, Huber W, Anders S. Moderated estimation of fold change and dispersion for RNA-seq data with DESeq2. Genome Biol. 2014;15:550.

**Supplementary Table S1** Clinical and pathological paremeters of prostate cancer cohort.

| **Clinical and pathological parameters** | | **n** | **%** |
| --- | --- | --- | --- |
| **Age** | ≤median (66) | 58 | 49.6 |
|  | >median (66) | 59 | 50.4 |
|  | Total | 117 |  |
| **T status** | pT2 | 60 | 53.6 |
|  | pT3 | 52 | 46.4 |
|  | Total | 112 |  |
| **Gleason score** | 3+4 | 35 | 31.3 |
|  | 4+3 | 44 | 39.3 |
|  | ≥8 | 33 | 29.5 |
|  | Total | 112 |  |
| **Pre-operative PSA** | <10 ng/ml | 80 | 68.4 |
|  | ≥10 ng/ml | 37 | 31.6 |
|  | Total | 117 |  |
| **Biochemical recurrence** | No | 71 | 67.0 |
|  | Yes | 35 | 33.0 |
|  | Total | 106 |  |
| **Metastasis** | No | 110 | 94.0 |
|  | Yes | 7 | 6.0 |
|  | Total | 117 |  |

**Supplementary Table S2** Antibody list used in the study.

| **Target** | | **Clone/Cat.#** | **Company** | **Dilution** | |
| --- | --- | --- | --- | --- | --- |
|  |  |  |  | **Immunofluorescence staining** | **Western blotting** |
| **Emerin** | | 4G5 | Novocastra | 1:500 | 1:2000 |
| **Emerin** | | PA529731 | ThermoFisher Sci. | 1:500 | 1:2000 |
| **Sec61B** | | ab244487 | abcam | 1:100 |  |
| **LBR (Lamin B receptor)** | | ab232731 | abcam | 1:200 |  |
| **cGAS** | | D1D3G | Cell Signaling | 1:200 |  |
| **SUN2** | | ab124916 | abcam | 1:100 |  |
| **LAP2alpha** | | 3A3 | Cell Signaling | 1:100 |  |
| **BAF-1** | | A-11 X | Santa Cruz Biotechnology | 1:100 |  |
| **γH2AX** | | 20E3 | Cell Signaling | 1:200 |  |
| **SUN1** | | ab124770 | abcam | 1:100 |  |
| **H3K27me3** | | C36B11 | Cell Signaling | 1:100 |  |
| **Lamin B1** | | ab16048 | abcam | 1:200 | 1:2000 |
| **p62** | | ab194721 | abcam | 1:200 |  |
| **Lamin A/C** | | ab215495 | abcam | 1:200 |  |
| **Lamin A/C** | | MA3-1000 | ThermoFisher Sci. | 1:500 | 1:2000 |
| **Nesprin 1** | | ab192234 | abcam | 1:100 |  |
| **SMC3** | | PA5-29131 | ThermoFisher Sci. | 1:200 | 1:1000 |
| **BRCA2** | | HPA026815 | Sigma-Aldrich |  | 1:1000 |
| **XRCC2** | | HPA065153 | Sigma-Aldrich |  | 1:1000 |
| **Pericentrin** | | ab28144 | abcam | 1:200 |  |
| **Pericentrin** | | ab270119 | abcam | 1:200 |  |
| **Paxilin** | | ab32084 | abcam | 1:200 |  |
| **α-Tubulin** | | T5168 | Sigma-Aldrich |  | 1:5000 |
| **β-actin** | | AC-74 | Sigma-Aldrich |  | 1:10000 |
| **Secoundary antibodies** | | | | | |
| **Goat anti-Rabbit IgG (H+L) Alexa Fluor™ 488** | | A11008 | ThermoFisher Sci. | 1:500 |  |
| **Goat anti-Rabbit IgG (H+L) Alexa Fluor™ 546** | | A11010 | ThermoFisher Sci. | 1:500 |  |
| **Goat anti-Rabbit IgG (H+L) Alexa Fluor™ 647** | | A21244 | ThermoFisher Sci. | 1:500 |  |
| **Goat anti-Mouse IgG (H+L) Alexa Fluor™ 488** | | A11001 | ThermoFisher Sci. | 1:500 |  |
| **Goat anti-Mouse IgG (H+L) Alexa Fluor™ 546** | | A11003 | ThermoFisher Sci. | 1:500 |  |
| **Goat anti-Mouse IgG (H+L) Alexa Fluor™ 647** | | A21235 | ThermoFisher Sci. | 1:500 |  |
| **AlexaFluor® 680-conjugated AffiniPure Goat Anti-Rabbit** | | 111-625-144 | Jackson ImmunoResearch | - | 1:12500 |
| **AlexaFluor® 790-conjugated AffiniPure Donkey Anti-Mouse** | | 715-655-150 | Jackson ImmunoResearch | - | 1:12500 |

**Supplementary Table S3** RNAi sequences used in the study

| **Target Gene Symbol** | **siRNA ID** | **Sense** | **Antisence** |
| --- | --- | --- | --- |
| **SMC3** | s17427 | GCCUAAGCAACGUAGCUUAtt | UAAGCUACGUUGCUUAGCat |
| **LMNA** | 144426 | GGAGCUGAAAGCGCGCAAUtt | AUUGCGCGCUUUCAGCUCCtt |
| **LMNB1** | 144054 | GCUCUUGCUACUGCACUUGtt | CAAGUGCAGUAGCAAGAGCtg |
| **BRCA2** | s2085 | GGAUUAUACAUAUUUCGCAtt | UGCGAAAUAUGUAUAAUCCag |
| **XRCC2** | s14945 | GGCUAGUUACAAUUCUUGAtt | uCAAGAAUUGUAACUAGCCgg |

**Supplementary Table S4** Differentially expressed genes among tumors with EMD-rich MN or EMD pauperized form NE

| **Symbol** | **Entrez** | **log2FC** | **p-val** |
| --- | --- | --- | --- |
| SRGN | 5552 | 0.87 | 0.002 |
| CXCR4 | 7852 | 0.67 | 0.009 |
| COL1A2 | 1278 | 2.04 | 0.009 |
| COL5A2 | 1290 | 0.47 | 0.009 |
| APOE | 348 | 1.51 | 0.011 |
| COL3A1 | 1281 | 0.98 | 0.012 |
| SPARC | 6678 | 0.96 | 0.013 |
| CTSK | 1513 | 0.43 | 0.013 |
| VIM | 7431 | 1.64 | 0.018 |
| LUM | 4060 | 0.91 | 0.019 |
| TGFB1 | 7040 | 0.29 | 0.020 |
| EPHB4 | 2050 | 0.43 | 0.021 |
| EMILIN1 | 11117 | 1.09 | 0.024 |
| GSN | 2934 | 1.17 | 0.027 |
| AEBP1 | 165 | 1.34 | 0.029 |
| ID4 | 3400 | 1.31 | 0.029 |
| IGFBP7 | 3490 | 1.39 | 0.031 |
| TGFBR2 | 7048 | 0.81 | 0.032 |
| IGFBP4 | 3487 | 1.12 | 0.032 |
| COL6A3 | 1293 | 0.85 | 0.033 |
| C1S | 716 | 0.84 | 0.033 |
| TCF4 | 6925 | 0.67 | 0.033 |
| CCL5 | 6352 | 0.72 | 0.035 |
| ANXA2P2 | 304 | 0.66 | 0.035 |
| SMAD3 | 4088 | 0.30 | 0.038 |
| COL1A1 | 1277 | 0.78 | 0.040 |
| SFRP1 | 6422 | 0.72 | 0.042 |
| MMP2 | 4313 | 0.77 | 0.042 |
| MAP2K4 | 6416 | -0.21 | 0.043 |
| COL18A1 | 80781 | 1.31 | 0.043 |
| THBS2 | 7058 | 0.31 | 0.044 |
| FN1 | 2335 | 1.28 | 0.045 |
| PIK3R1 | 5295 | 0.27 | 0.047 |
| CCDC80 | 151887 | 0.50 | 0.050 |

**Supplementary Table S5** The top 9 up-regulated genes in both datasets – RNA sequencing of PC-3 (control vs EMD-KO) and Nanostring analysis of g tumors with EMD-rich MN or EMD pauperized form NE.

| **external_gene_name** | **log2FC RNAseq** | **p-val RNAseq** | **log2FC Nanostring** | **p-val Nanostring** |
| --- | --- | --- | --- | --- |
| CXCR4 | 3.31 | 1.50127E-08 | 0.67 | 0.0085459 |
| APOE | 1.07 | 0.00182509 | 1.51 | 0.0105515 |
| SPARC | 0.62 | 0.041990448 | 0.96 | 0.0129548 |
| VIM | 0.98 | 1.36492E-09 | 1.64 | 0.0178852 |
| GSN | 1.01 | 7.49137E-12 | 1.17 | 0.0271238 |
| ANXA2P2 | 0.66 | 0.001324555 | 0.66 | 0.0351556 |
| SFRP1 | 1.17 | 0.000457857 | 0.72 | 0.0415546 |
| COL18A1 | 0.50 | 2.4982E-05 | 1.31 | 0.0429317 |
| FN1 | 0.69 | 7.89609E-05 | 1.28 | 0.0449681 |

**Supplementary Movie 1**

Live cell imaging of mitosis of PC-3 cells stably expressing EMD-EGFP construct, treated with IRCF-193 inhibitor. The cell division is associated with chromatin bridge formation and EMD-rich MN formation after its resolution. Scale bar 10 μm, time in minutes. EMD-GFP is color coded.

**Supplementary Movie 2**

Live cell imaging of mitosis of PC-3 cells stably expressing EMD-EGFP construct, treated with IRCF-193 inhibitor. The cell division is associated with chromatin bridge formation and EMD-rich MN formation during its resolution. Scale bar 10 μm, time in minutes. EMD-GFP is color coded.

**Supplementary Movie 3**

Live cell imaging of PC-3 cells stably expressing EMD-EGFP construct, treated with IRCF-193 inhibitor. The cell division ends up in bi-nucleated cell formation. Scale bar 10 μm, time in minutes. EMD-GFP is color coded.

**Supplementary Movie 4**

iFRAP (inverse florescence recovery after photobleaching) of nuclear envelope in of PC-3 cells stably expressing EMD-EGFP construct. Scale bar 10 μm, time in seconds.

**Supplementary Movie 5**

iFRAP (inverse florescence recovery after photobleaching) of EMD-rich MN in of PC-3 cells stably expressing EMD-EGFP construct. Scale bar 10 μm, time in seconds.

**Supplementary Movie 6**

Comparison of 2D migration properties of PC-3 cells stably expressing EMD-EGFP construct in cell with nuclear envelope localization of EMD and a cell with EMD-rich MN. Scale bar 10 μm, time in minutes.
